# Environmental determinants of suitable habitat and the prediction of a southern shift in the future distribution of spiny lobsters, genus *Jasus*

**DOI:** 10.1101/2023.04.28.538751

**Authors:** Jason H. E. Tepker, Jan M. Strugnell, Catarina N. S. Silva

## Abstract

Climate change is altering environmental conditions which is affecting species habitats globally. As a result, many species are shifting their habitat ranges poleward to ensure that they remain within their optimal thermal range. On average, marine species have shifted their ranges poleward by approximately 72km to track their optimal thermal conditions compared to 17km for terrestrial species. These range shifts are pushing some species out of their currently fished areas. This will require nations and fishery companies to predict the most likely areas their target species could move to and obtain permits to fish in those new areas. Spiny lobsters (genus *Jasus*) are likely to shift their ranges poleward as they are distributed within a relatively tight latitudinal band, but there is limited information about the extent of any potential range shifts. The aims of this study were to identify the environmental variables that characterise the current habitat locations of lobsters within the genus *Jasus*, and to predict their potential distribution by modelling future suitable habitat under the RCP45, RCP60, and RCP85 climate scenarios using MaxEnt. There were 16 environmental variables used for modelling suitable habitat for the present (2000-2014), while only four environmental variables were available for modelling in two future time periods (2040-2050 and 2090-2100). There was a predicted overall southern shift in suitable habitat locations for all species. The most important environmental variable identified for species distributed along continental shelves (*J. edwardsii* and *J. lalandii*) was benthic temperature. Benthic nutrients (silicate, nitrate, and phosphate) were the most important variables for species distributed around islands and on seamounts (*J. paulensis, J. frontalis,* and *J. tristani*). Approximately 90% of *Jasus* lobsters’ present range contained highly suitable habitat locations. The percent of highly suitable locations under the RCP45 and RCP60 scenarios were higher than the present percentages for each species, while under the RCP85 scenario, there was a decrease of highly suitable habitat for most species in 2040-2050 period, while for the 2090- 2100 period, there was an increase in the percent of highly suitable habitats. This study provides evidence that *Jasus* populations might become more abundant in the southern extents of their current range as they track their optimum habitat conditions.

## 1.0. Introduction

Over the past century, the effects of climate change and other human impacts on the oceans have affected the habitats of marine species. These impacts include habitat deterioration (Provost et al. 2017; Gravinese 2018), changes in ocean circulation affecting larval dispersion (Cetina-Heredia et al. 2015), and ocean warming (Doney et al. 2012). These and other impacts are forcing species to shift their ranges in order to find more suitable habitat locations (Tyberghein et al. 2012; Carlos-Júnior et al. 2015). Many marine species are shifting their ranges poleward and to deeper habitats in order to remain within their optimal thermal window (Fitzgibbon et al. 2014; Oliver & Holbrook 2014; Pecl et al. 2017). Marine species have been reported to be migrating poleward at approximately 72 km per decade in order to find new suitable habitats, while terrestrial species are moving more slowly at around 17 km per decade (Pecl et al. 2017). However, each species will be required to move different distances in order to find suitable habitat where the environmental conditions fall within their tolerance levels (Pecl et al. 2017).

Different environmental variables can determine the suitability of habitats (McHenry et al. 2019). Marine lobsters are influenced by, and have optimal ranges, for environmental variables such as water temperature, dissolved oxygen content, pH, salinity, and depth (Fitzgibbon et al. 2014; Milton et al. 2014; Bradford et al. 2015). They have a thermal window within which physiological performance is optimal (Crear & Forteath 2000; Funes-Rodriguez et al. 2015; Provost et al. 2017). Fluctuations to water temperature affect the amount of dissolved oxygen available, a parameter which is vitally important for survival (Knapp 2015; Senevirathna et al. 2017). Temperature changes also affect salinity which in turn influences physiological processes such as growth and development (Bermudes & Ritar 2005; Vidya & Joseph 2012). In addition, warming temperatures caused by rising carbon dioxide levels are decreasing the pH of the oceans which negatively affects calcite concentrations (Hinojosa et al. 2017). Fluctuations in pH levels require species to allocate more energy toward maintaining their internal processes while reducing the amount of energy available for other processes such as growth and reproduction (Verghese et al. 2007; Knapp et al. 2016; Gravinese 2018).

Marine lobsters, from the genus *Jasus,* are distributed within a relatively narrow latitudinal range across the southern hemisphere (Figure 1). *J. edwardsii* is broadly distributed across the southern continental shelf regions of Australia and is also distributed around New Zealand (MacDiarmid et al. 2011b). *J. lalandii* inhabits continental shelf regions along the south and west coasts of South Africa (Cockcroft et al. 2011a). While *J. edwardsii* and *J. lalandii* are geographically distant, they share a common environment (continental shelf) and ancestor despite *J. lalandii* being geographically closer to *J. paulensis* and *J. tristani* (Silva et al. 2021a). *J. paulensis* has a small range distribution located in the southern Indian ocean around two remote islands (St. Paul and Amsterdam) and along nearby seamounts (Cockcroft et al. 2011b). *J. tristani* has a small range located on the Vema Seamount, and around Gough Island in the South Atlantic Ocean (Cockcroft et al. 2011c). *J. paulensis* and *J. tristani* share a common ancestor and habitat environment (island and seamount habitat) compared to the other four species (Silva et al. 2021a). Genetic differentiation between *J. paulensis* and *J. tristani* is low but significant, and patterns of connectivity between these species changed over time in accordance with climatic fluctuations (Silva et al. 2021b). *J. frontalis* inhabits areas around the Islas Desventuradas and the Juan Fernandez Archipelago off the coast of Chile (Wahle et al. 2011). *J. caveorum* is located along the Foundation Seamount chain (35° S, 120° W) located in the southeastern Pacific Ocean (MacDiarmid et al. 2011a) (Figure 1). *J. frontalis* and *J. caveorum* share a common ancestor and are more genetically differentiated from the previously mentioned species possibly due to their isolated locations and oceanic processes limiting larval dispersion in the Eastern Pacific Ocean (Silva et al. 2021a).

**Figure 1.**
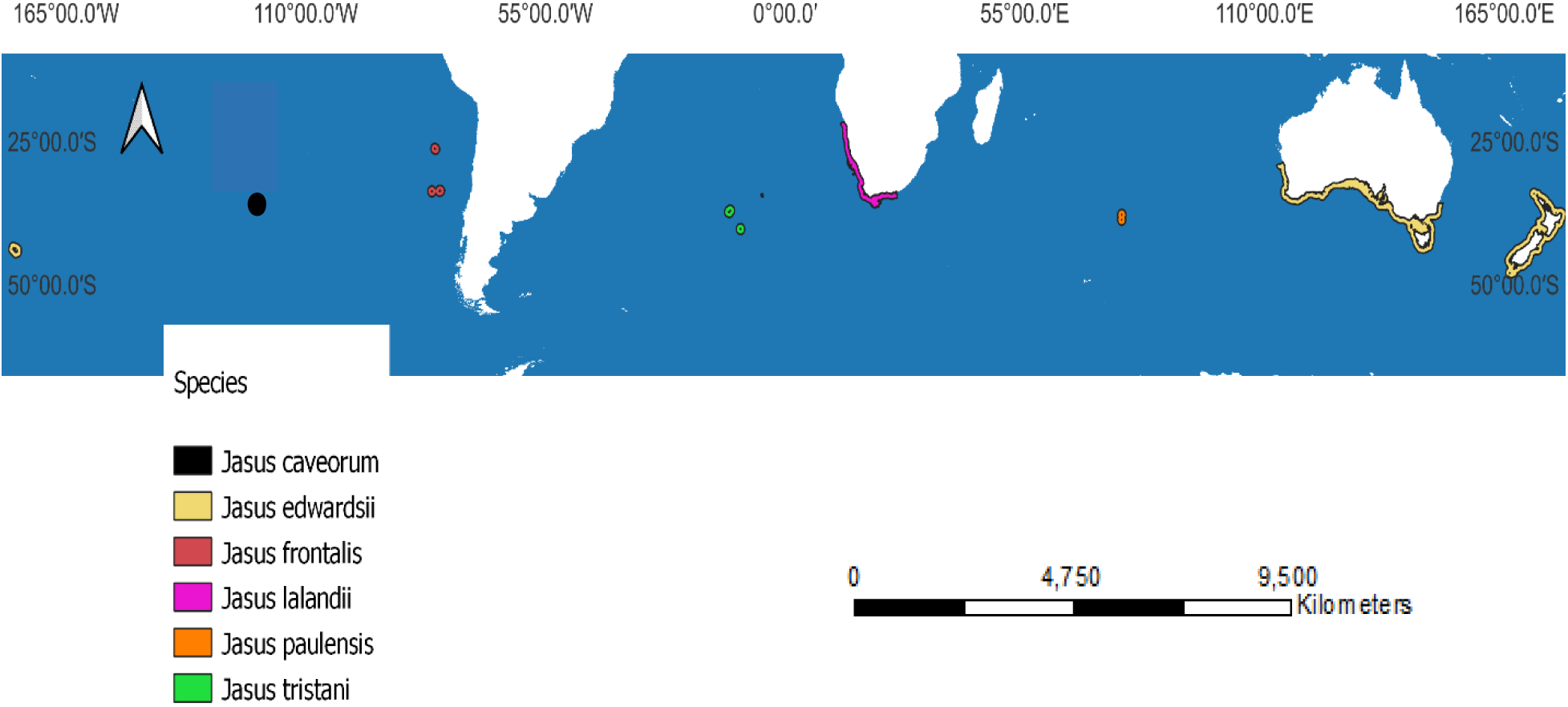
Estimated spatial distributions for the six *Jasus* species current habitat ranges. Data downloaded from GBIF (GBIF.org (accessed on 17 October 2022) GBIF Occurrence Download https://doi.org/10.15468/dl.tqj6wd). The map is using the CGS WGS 1984 coordinate system.

Most *Jasus* lobster species support national and commercial fisheries, with the exception of *J. caveorum* (Jeffs et al. 2013). The largest commercial fishery is for *J. edwardsii* which generates approximately $450 million USD per year (Linnane & Crosthwaite 2009; Hinojosa et al. 2015). The fishery for *J. lalandii* has dropped from an annual catch of 18,000 tonnes in the early 1950’s to approximately 2,000 tonnes currently (Johnston and Butterworth 2005). However, in the 1980’s and 1990’s, *J. lalandii* was discovered in high densities (50 to 100 fold increase) along the southwest coast of South Africa where it was previously rare to find them (Haley et al. 2011). As a result, it remains a valuable fishery for South Africa worth approximately $40 million USD (Johnston and Butterworth 2005). The fisheries for *J. paulensis, J. tristani,* and *J. frontalis* are significantly smaller with annual landings of approximately 390 tonnes, 380 tonnes, and 58 tonnes respectively (Jeffs et al. 2013). While *J. caveorum* does not support a consistent fishery, there are estimates that approximately 20 tonnes have been harvested since their discovery in 1995 (Jeffs et al. 2013). *J. frontalis, J. paulensis, J. tristani,* and *J. caveorum* are distributed in more remote locations (around islands and along seamounts) which could deter some fisheries from venturing into those more isolated and unpredictable oceanic regions. All of the *Jasus* fisheries have seen decreases in annual landings in recent years (Johnston and Butterworth 2005; Jeffs et al. 2013), and any changes to their habitats will likely have a profound impact on the fisheries.

Species within the genus *Jasus* have one of the longest known larval stages which can last for up to two years (in *J. edwardsii*), and therefore they have the potential for widespread dispersal (Bradford et al. 2015; Hinojosa et al. 2016; Chiswell & Booth 2017). Lobster larvae are thought to be transported by oceanic processes potentially hundreds of kilometres away from their origin (Chiswell & Booth 1999; Young et al. 2016; Hinojosa et al. 2017). Once the larvae metamorphose into pueruli, they are able to return to their preferred habitats through active swimming using habitat cues (chemical, audio, etc.) (Hinojosa et al. 2018). Adults mainly inhabit kelp and rocky reef habitats located within the upper 200m of the water column, but individuals of some species have been found at depths of 600m (Booth 2002; Jeffs et al. 2013; Hesse et al. 2015). They inhabit structurally complex habitat as juveniles require holes and crevices to hide from predators (Booth 2002; Hesse et al. 2015, 2016).

Despite their long larval duration and potential for long distance dispersal, *Jasus* lobsters are distributed within a narrow latitudinal band. This raises the question about what are the environmental variables influencing the relatively limited distribution of *Jasus* lobsters. The aims of this paper were to identify the environmental variables that are determining the spatial distribution for *Jasus* lobsters and to predict future suitable habitat under three climate change scenarios. First, the importance of each environmental variable was determined, and that information was then used to predict the suitability of areas within and around the current range for each *Jasus* lobster species. Then, future climate projection data was used to simulate how three climate scenarios (RCP45, RCP60, and RCP85) might affect suitable habitats for *Jasus* lobsters in two future periods (2040-2050 and 2090-2100).

### 2.0. Methods

## 2.1. Environmental variables

The bathymetry layer (SRTM30 Plus) is a global raster layer downloaded from the University of California San Diego’s oceanography department website (https://topex.ucsd.edu/WWW_html/srtm30_plus.html). It uses a satellite-gravity model to obtain a higher grid cell resolution and represents the ocean’s depth profile (Becker et al. 2009). It has a spatial resolution of 30 arc seconds corresponding to a raster grid cell covering a one-kilometre area at the equator (Becker et al. 2009).

Additional environmental variables are available for download from the Bio-ORACLE marine datasets website (http://www.bio-oracle.org/) (Table 1). Bio-ORACLE’s global marine datasets are composed of numerous environmental variables with high spatial resolutions at the ocean surface and benthic regions (Tyberghein et al. 2012; Assis et al. 2017). There are 23 environmental variables available of which 15 were downloaded for this study as they are more biologically relevant to *Jasus* lobsters compared to the remaining eight. All environmental variables have spatial resolutions of five arcminutes which equates to the raster cells covering a 9.2-kilometre area at the equator (Tyberghein et al. 2012; Assis et al. 2017). The benthic data was cross-validated with *in-situ* data, and represented important environmental aspects in marine species distributions to be used for bioclimatic modelling (Tyberghein et al. 2012; Assis et al. 2017). The downloaded environmental variables for the present time period (2000-2014) use mean cell values, while the benthic layers represent the mean depth and use mean cell values. Surface layers are available for all 15 environmental variables, while benthic layers are not available for calcite concentration, diffuse attenuation, photosynthetic available radiation, and pH (Table 1).

**Table 1.** Marine environmental variable raster layers downloaded from the Bio-ORACLE website (http://www.bio-oracle.org/). The abbreviations in the brackets correspond to how each variable is named for use in the modelling package MaxEnt. The category headings of surface and benthic layer indicate if each layer is available for that depth. The unit m represents metres. The unit mol represents a mole. The unit umol represents a micro mole. The unit km represent kilometres. The unit E represents energy. The unit g represents grams.

| Variable | Surface Layer | Benthic Layer | Layers Available in Future Periods | Cell Resolution (at equator) | Units of measure |
| --- | --- | --- | --- | --- | --- |
| Calcite (calcite_clip) | Yes | No | No | 9.2 km | mol/m<sup>3</sup> |
| Chlorophyll | Yes | Yes | No | 9.2 km | mg/m<sup>3</sup> |
| Current Velocity (current_clip) | Yes | Yes | Yes | 9.2 km | m/s |
| Dissolved Oxygen Content (oxygen_clip) | Yes | Yes | No | 9.2 km | mol/m<sup>3</sup> |
| Diffuse Attenuation (diffuse) | Yes | No | No | 9.2 km | m |
| Iron | Yes | Yes | No | 9.2 km | umol/m<sup>3</sup> |
| Nitrate | Yes | Yes | No | 9.2 km | mol/m<sup>3</sup> |
| Phosphate | Yes | Yes | No | 9.2 km | mol/m<sup>3</sup> |
| Photosynthetic Available Radiation (PAR) | Yes | No | No | 9.2 km | E/m<sup>2</sup>/day |
| Phytoplankton (phyto_clip) | Yes | Yes | No | 9.2 km | umol/m<sup>3</sup> |
| Primary Productivity (primary) | Yes | Yes | No | 9.2 km | g/m<sup>3</sup>/day |
| **pH (pH_clip)** | Yes | No | No | 9.2 km | Unit less (measures difference from average) |
| **Salinity (salinity_clip)** | Yes | Yes | Yes | 9.2 km | PSS (practical salinity scale) |
| **Silicate** | Yes | Yes | No | 9.2 km | mol/m<sup>3</sup> |
| **Temperature (temp_clip)** | Yes | Yes | Yes | 9.2 km | degrees Celsius |

Bio-ORACLE also provides climate projection data for two future time periods: 2040- 2050 and 2090-2100 (Tyberghein et al. 2012; Assis et al. 2017). Those layers represent the RCP45, RCP60, and RCP85 predictions which correspond to three different climate change scenarios (IPCC Synthesis Report Box 2.2: https://ar5-syr.ipcc.ch/topic_futurechanges.php). The environmental variables for these future time periods use mean cell values for the surface and the benthic layers. However, there are only three biologically relevant environmental variables with climate data available for these two future periods: current velocity, salinity, and temperature (Table 1).

## 2.2. Data preparation

The environmental variables were loaded into the ArcGIS software package ArcMap 10.6 (https://www.esri.com/en-us/arcgis/products/arcgis-desktop/overview) as TIF raster files. All Bio-ORACLE environmental layers were re-sampled to a spatial resolution of one-kilometre using bilinear interpolation, and maintained their CGS WGS 1984 coordinate system to match the bathymetry layer.

Data from all the environmental layers was extracted using an ArcMap custom-made shapefile that covered an area which included all the species current habitat ranges and surrounding areas. This was necessary because all the environmental layers were on a global scale and extracting the data over a smaller range decreased the modelling time substantially. The extents (in degrees) of the shapefile used to extract the data were -179.9167, -65.9167: 179.833, -9.667. Also, location shapefiles were created for each species habitat range from data downloaded from GBIF (GBIF.org (accessed on 17 October 2022) GBIF Occurrence Download https://doi.org/10.15468/dl.tqj6wd). Occurrence data for each species was also downloaded from GBIF. All environmental variable layers were converted to the ASC file type for modelling.

## 2.3. Environment modelling

The extracted environmental variable layers along with the individual species and genus occurrence points were loaded into the modelling program MaxEnt (version 3.4.1). MaxEnt is a maximum entropy niche modelling package which uses presence-only data (Phillips et al. 2006; Phillips & Dudik 2008). MaxEnt uses a machine-learning approach to determine the probability of importance for each environmental variable within the target species distribution, and then uses that information to find other potential suitable habitat areas (Phillips et al. 2006; Phillips & Dudik 2008). MaxEnt simulations were run for the genus and for each species to allow for comparisons at the individual species level and between species.

The MaxEnt settings used were: maximum iterations = 800, maximum number of background points = 10,000 (default setting), number of replicates = 3. There were three replicates to represent three different model runs to ensure that the most important environmental variable was the same for each run. The maximum iterations was increased from the default setting of 500 to 800 to ensure that maps were produced (Phillips et al. 2006; Phillips & Dudik 2008). The number of occurrence points used for training and testing varied for each species and for the genus, and was based on the total number of unique occurrence points within the original file (Table 2). Duplicate occurrence points were removed, and randomly selected occurrence points were used for statistical testing on each run. The results were produced using a logistic algorithm which creates maps using a scale ranging from 0 (not suitable locations represented as light blue areas) to 1 (highly suitable locations represented as red areas). An additional jackknife area under the curve (AUC) test was used to determine the importance of each environmental variable in the model. This test estimated the effect that each individual environmental variable had on the overall performance of the model by only using each variable independently, and also by excluding only that variable. Maps were created using QGIS 3.26.3 (https://www.qgis.org/en/site/) to show the suitability of areas.

**Table 2.**
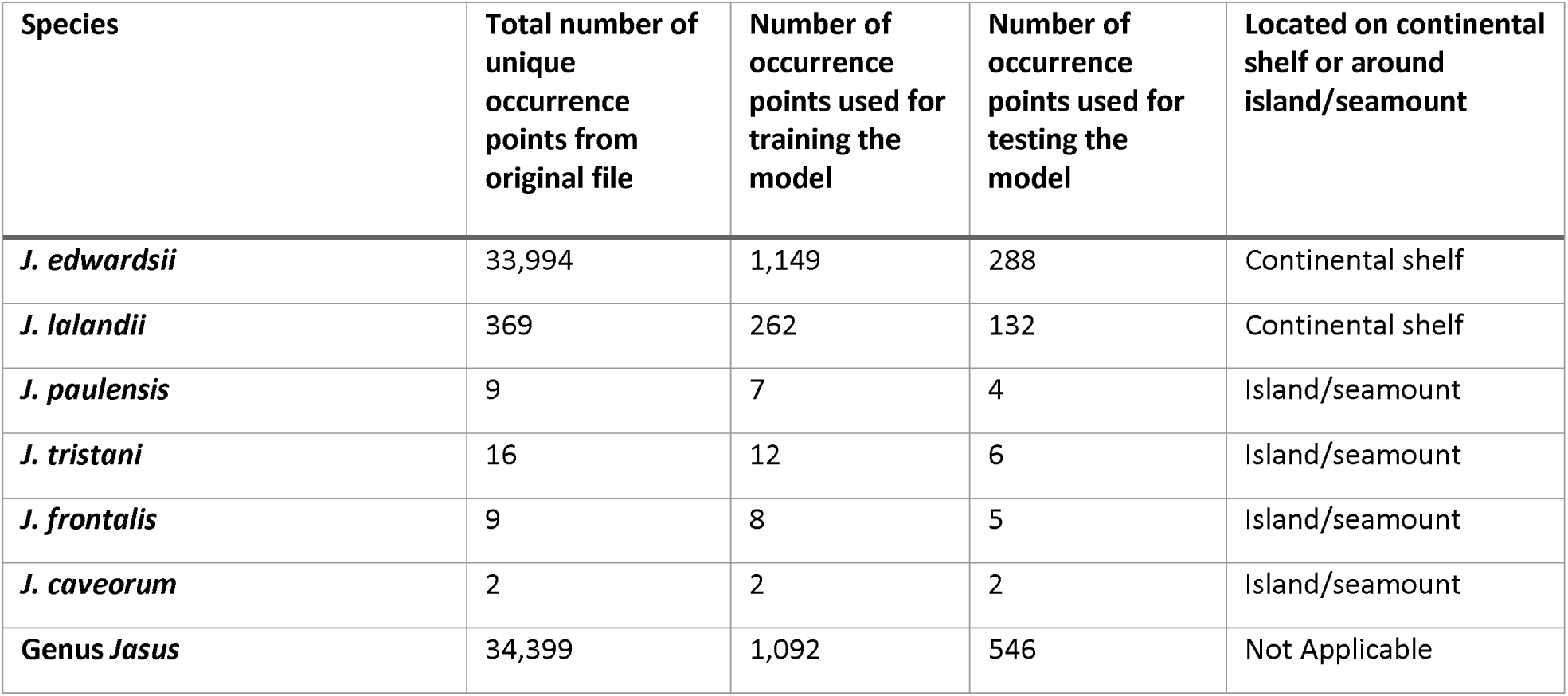
Total number of unique occurrence points, number of occurrence points used for model training and testing for *Jasus* lobsters downloaded from GBIF (GBIF.org (accessed on 17 October 2022) GBIF Occurrence Download https://doi.org/10.15468/dl.tqj6wd). The location of each species habitat range is included.

| Species | Total number of unique occurrence points from original file | Number of occurrence points used for training the model | Number of occurrence points used for testing the model | Located on continental shelf or around island/seamount |
| --- | --- | --- | --- | --- |
| *J. edwardsii* | 33,994 | 1,149 | 288 | Continental shelf |
| *J. lalandii* | 369 | 262 | 132 | Continental shelf |
| *J. paulensis* | 9 | 7 | 4 | Island/seamount |
| *J. tristani* | 16 | 12 | 6 | Island/seamount |
| *J. frontalis* | 9 | 8 | 5 | Island/seamount |
| *J. caveorum* | 2 | 2 | 2 | Island/seamount |
| Genus *Jasus* | 34,399 | 1,092 | 546 | Not Applicable |

## 2.4. Future climate modelling

For future projections, additional ASC files were included in MaxEnt to run simulations for the two future periods of 2040-2050 and 2090-2100 for the three climate scenarios of RCP45, RCP60, and RCP85. Given the software’s limitation of only being able to use the environmental variables that had both future climate projection data and present data, the only Bio-ORACLE environmental variable layers used were: surface and benthic layers for current velocity and temperature, and the benthic salinity layer. It was assumed that the bathymetry would not change between the present and the two future periods, and it was also included.

### 3.0. Results

## 3.1. Habitat modelling for the present period

The most important environmental variable in determining suitable habitat locations for *Jasus* lobsters was benthic temperature (Figure 2). Benthic temperature had the highest gain indicating that it is the main driver of suitable conditions in the model (Figure 2). The other highly important environmental variables were: primary productivity, phytoplankton concentration, and bathymetry (Table 3).

**Figure 2.**
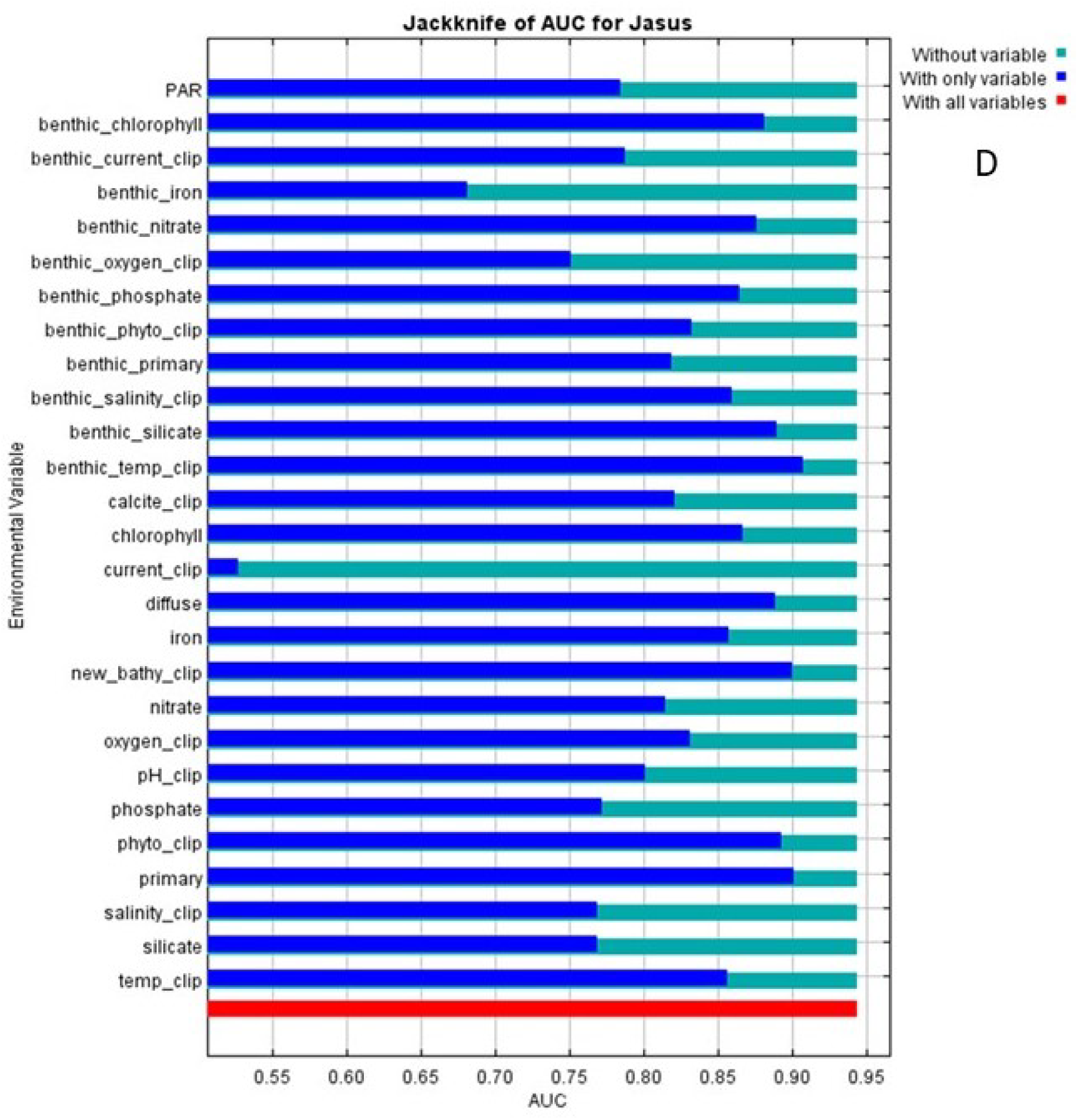
Area under the curve (AUC) jackknife results from MaxEnt simulations for genus *Jasus* for the present (2000-2014) indicating the importance of each environmental variable to the model when using only that variable (dark blue) or when excluding that variable (light blue).

**Table 3.**
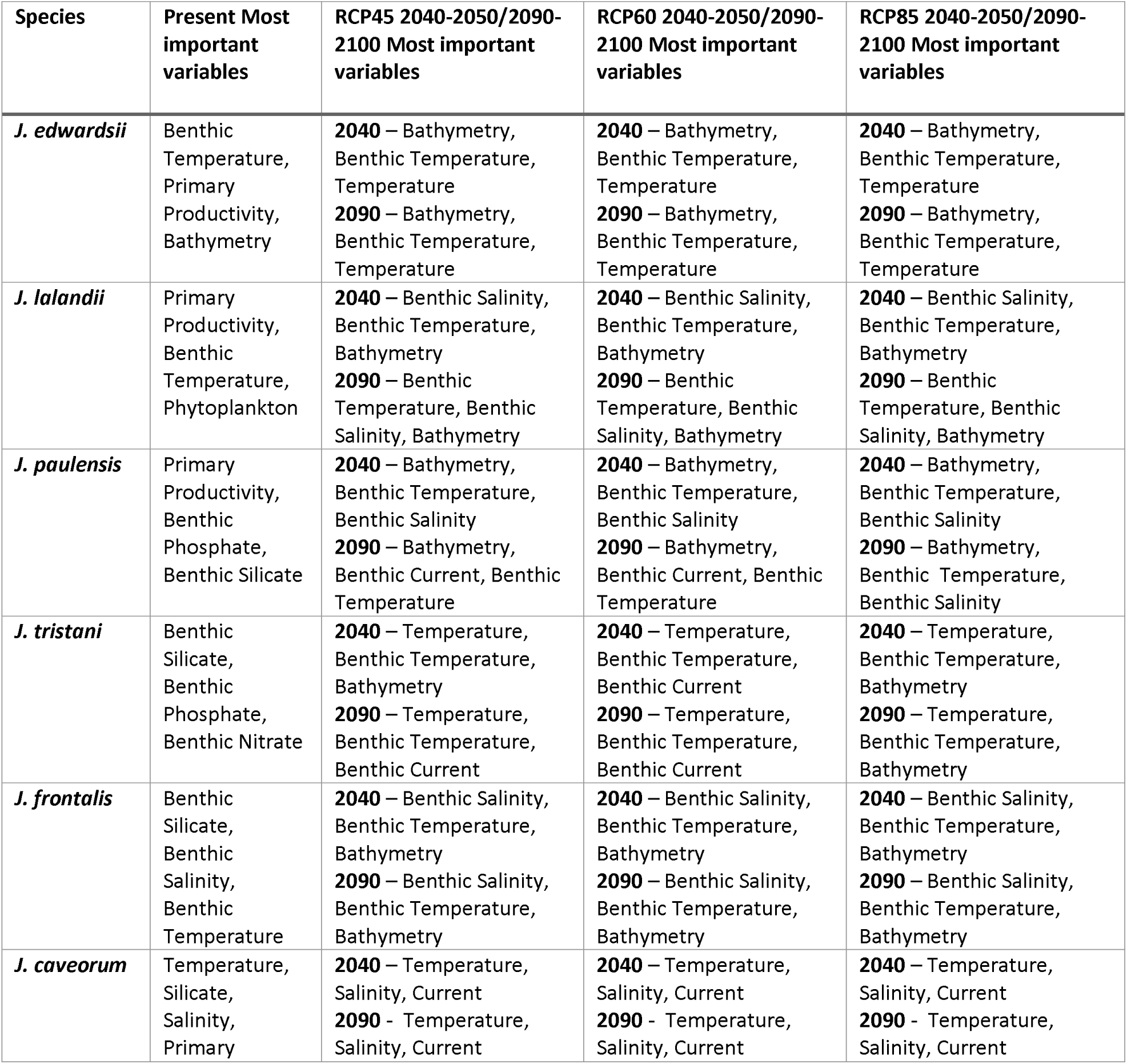
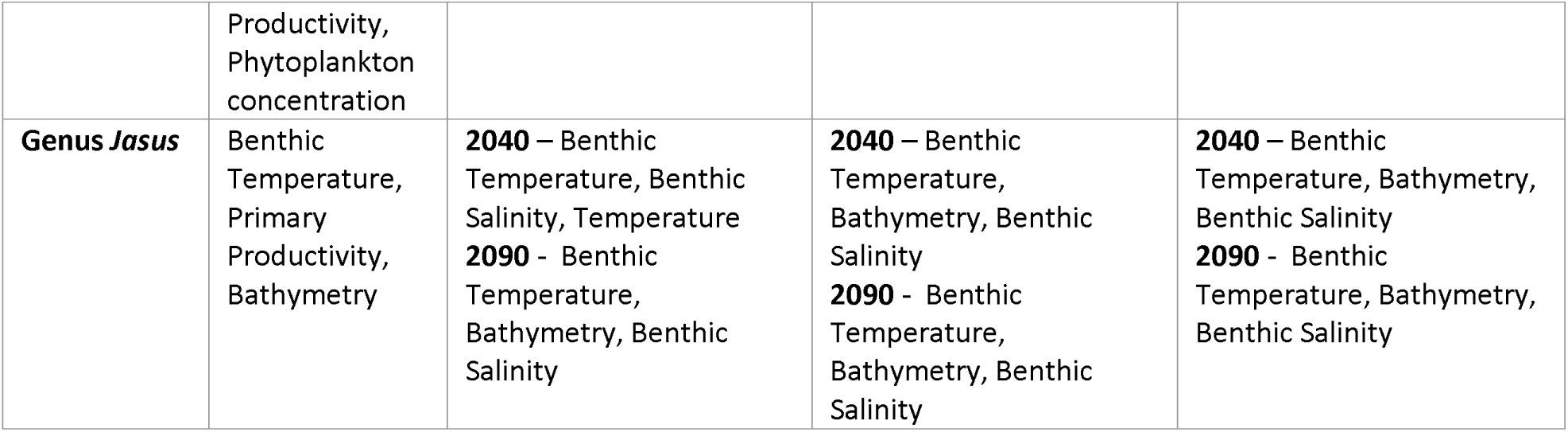
Summary of MaxEnt jackknife area under the curve (AUC) results showing the most important environmental variables for the *Jasus* genus and for each species for the present and for the two future periods (2040-2050 and 2090-2100) under climate scenarios RCP45, RCP60, and RCP85. More detailed results for genus *Jasus* and for each species are in the Supplementary Material Figures 2-8.

The most important environmental variable for each species differed, but there were similarities between species living in similar environment regions. For *J. edwardsii* and *J. lalandii* which inhabit continental shelf regions, benthic temperature and primary productivity were two of the most important environmental variables (Table 3). For *J. paulensis*, *J. frontalis*, and *J. tristani*, the most important variables were benthic nutrients (phosphate, silicate, and nitrate) indicating that species located on islands or seamounts may be more influenced by the availability of these nutrients. For *J. caveorum*, the most important variables were a mix of various surface and benthic variables (Table 3). However, results for *J. paulensis*, *J. frontalis*, *J. tristani,* and *J. caveorum* need to be interpreted with caution as there were limited numbers of occurrence points available for modelling.

Approximately 90% of the current habitat range for *Jasus* lobsters contained highly suitable conditions (Table 4). The percent of highly suitable locations within each individual species habitat varied. For *J. edwardsii,* its range contained 88% highly suitable locations. For *J. lalandii, J. frontalis,* and *J. paulensis*, the percentages were lower at 65%, 52%, and 45% respectively. The current ranges for *J. caveorum* and *J. tristani* contained 0% highly suitable habitat (Table 4). This was due to their low number of occurrence points and their small habitat ranges resulting in the same values for environmental variables being used with every occurrence point during modelling.

**Table 4.** Percent of highly suitable habitat locations within each species range and for the genus *Jasus* based on MaxEnt results for the present (2000-2014) and two future periods (2040-2050 and 2090-2100) under three climate scenarios (RCP45, RCP60, and RCP85). More detailed information in the Supplementary Material Figure 1.

| Species | Percent of highly<br>suitable locations in<br>the present | Percent of highly<br>suitable locations<br>in RCP45 | Percent of highly<br>suitable locations<br>in RCP60 | Percent of highly<br>suitable locations<br>in RCP85 |
| --- | --- | --- | --- | --- |
| *J. edwardsii* | 88% | 2040 – 90%<br>2090 – 95% | 2040 – 95%<br>2090 – 95% | 2040 – 79%<br>2090 – 85% |
| *J. lalandii* | 65% | 2040 – 75%<br>2090 – 80% | 2040 – 75%<br>2090 – 80% | 2040 – 80%<br>2090 – 90% |
| *J. paulensis* | 45% | 2040 – 71%<br>2090 – 61% | 2040 – 71%<br>2090 – 61% | 2040 – 27%<br>2090 – 71% |
| *J. tristani* | 0% | 2040 – 50%<br>2090 – 50% | 2040 – 50%<br>2090 – 60% | 2040 – 31%<br>2090 – 40% |
| *J. frontalis* | 52% | 2040 – 21%<br>2090 – 21% | 2040 – 21%<br>2090 – 21% | 2040 – 18%<br>2090 – 30% |
| *J. caveorum* | 0% | 2040 – 0%<br>2090 – 0% | 2040 – 0%<br>2090 – 3% | 2040 – 0%<br>2090 – 0% |
| **Genus *Jasus*** | 88% | 2040 – 92%<br>2090 – 95% | 2040 – 95%<br>2090 – 95% | 2040 – 78%<br>2090 – 85% |

Some highly suitable locations occurring outside of any species current ranges were found along the east coasts of Australia and South Africa, and off the east coast of New Zealand (Figure 3B, 3C). Also, there were large areas around several of the current habitat ranges of *Jasus* lobsters that contained slightly suitable locations. Overall, most of the areas outside of *Jasus* habitat ranges contain zero suitable conditions, except for locations within -20°S and -50°S (Figure 3).

**Figure 3.**
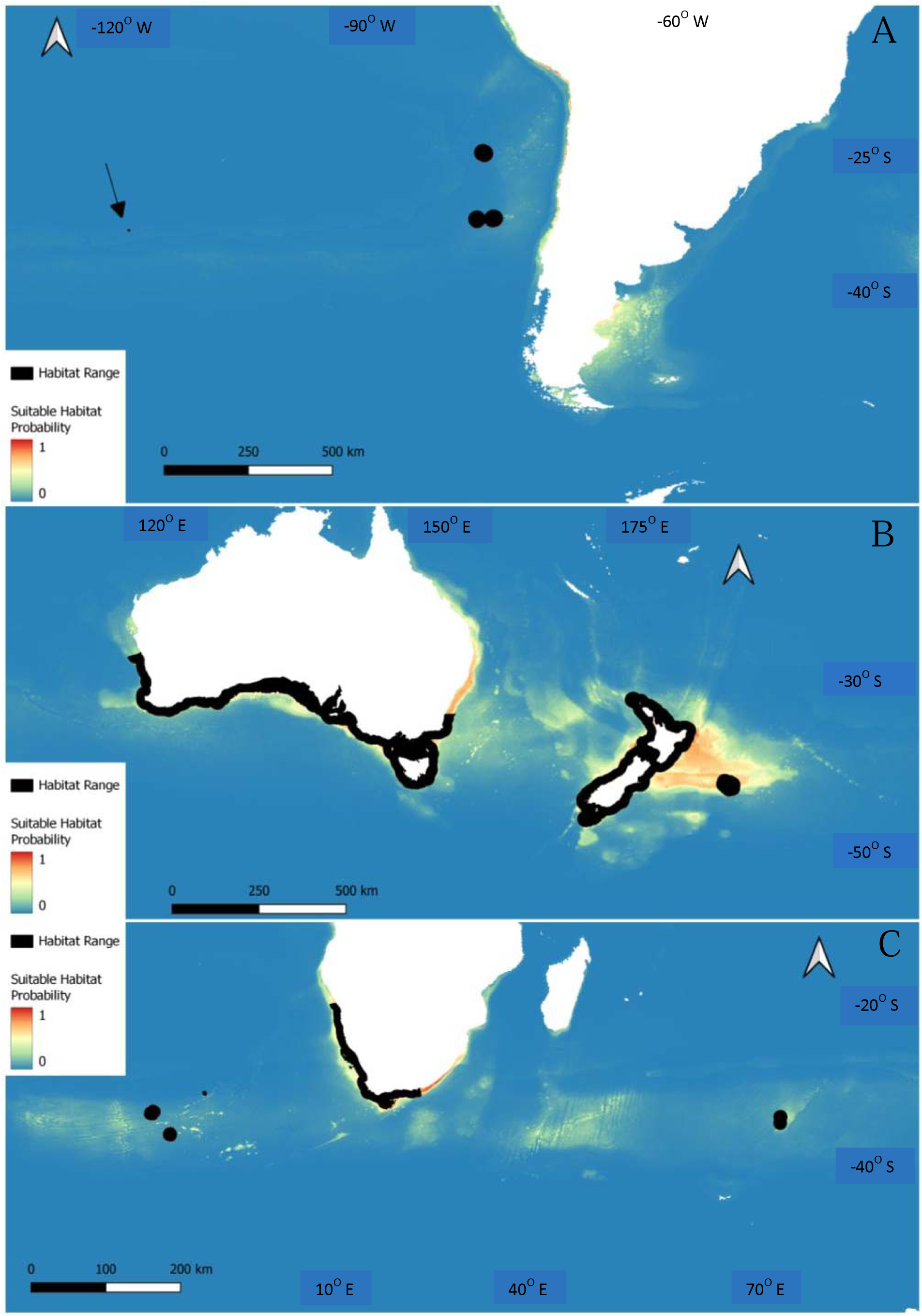
MaxEnt habitat suitability maps for the present period (2000-2014) for *Jasus* lobsters. Map A shows the habitat suitability results and ranges for *J. frontalis* which are the black dots off the west coast of South America, and *J. caveorum* which is the black dot beneath the black arrow to the west of *J. frontalis*. Map B shows the habitat suitability results and range for *J. edwardsii* around southern Australia and New Zealand. Map C shows the habitat suitability results and ranges for *J. lalandii* around South Africa, *J. paulensis* which are the black dots to the east of *J. lalandii*, and *J. tristani* which are the black dots to the west of *J. lalandii*.

## 3.2. Habitat modelling for climate scenario RCP45

Benthic temperature was consistently one of the most important environmental variables for all species, except for *J. caveorum* (Table 3). Modelling for the 2040-2050 and 2090-2100 time periods under the RCP45 scenario mainly showed an increase of highly suitable habitat locations compared to the present period (Table 4). Most species ranges contained highly suitable locations covering at least 50% of the area, except for *J. frontalis* and *J. caveorum* (Table 4). For most species, the period of 2040-2050 had a slightly lower percent of highly suitable locations compared to 2090-2100. However, *J. caveorum* remained the only species where its habitat range contained zero highly suitable locations (Table 4).

There were new highly suitable locations found along the west coast of South Africa, off the east coast of South America, to the southeast of South Africa, and off both the west and east coasts of New Zealand for 2040-2050 (Figure 4A-C). There were fewer highly suitable locations outside of the current *Jasus* habitat range in 2090-2100. Those occurred in a narrow band off the east coast of New Zealand, to the southeast of South Africa, and in a large area off the east coast of South America (Figure 4D-F). Since only four environmental variables were used for these time periods, the results should be interpreted carefully. However, it does suggest that as climate conditions change under the RCP45 scenario, suitable habitat locations will move south (Figure 4).

**Figure 4.**
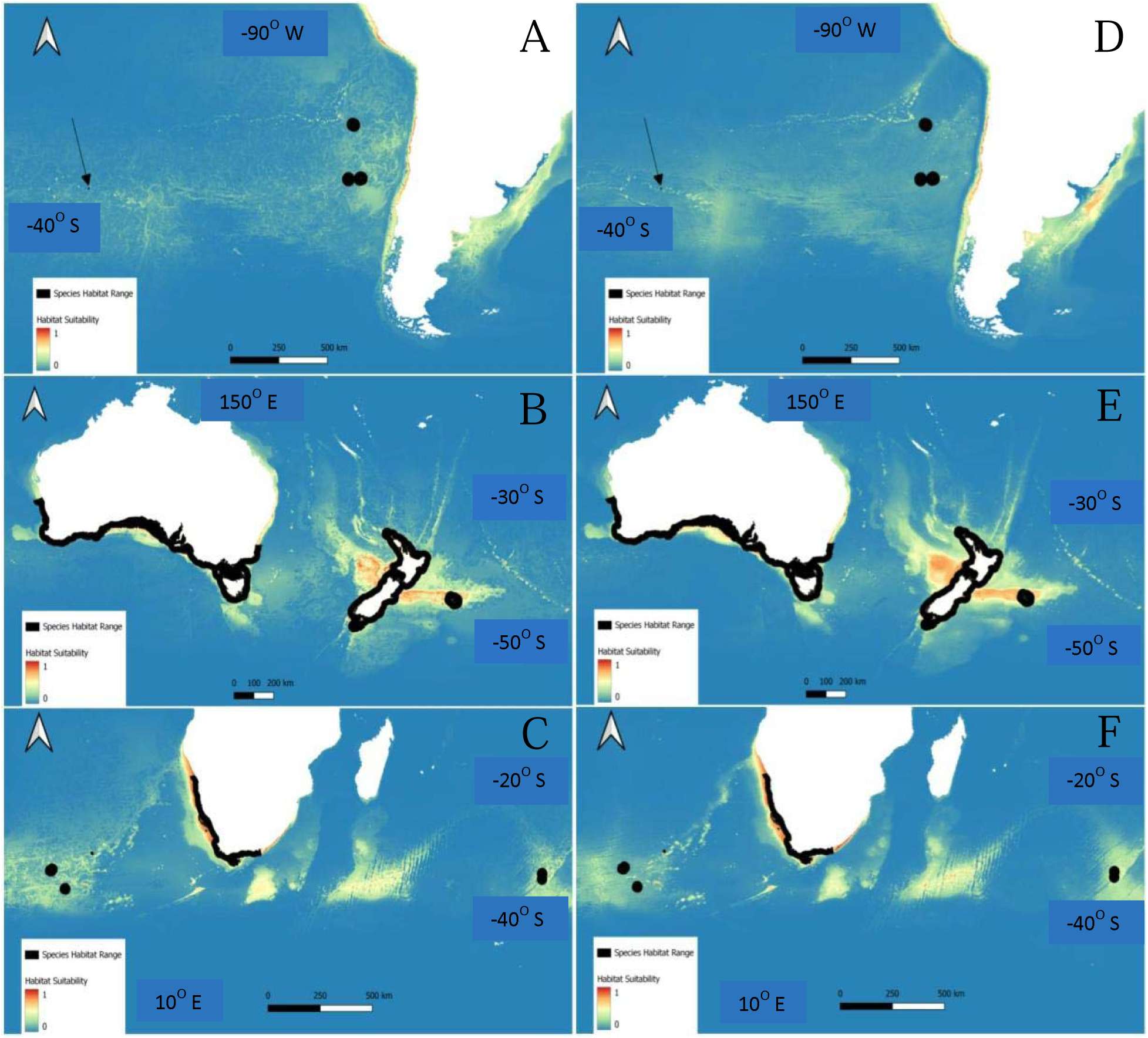
*Jasus* lobster MaxEnt habitat suitability maps for climate scenario RCP45 2040-2050 (A-C) and 2090-2100 (D-F) future periods. Maps A and D show the habitat suitability results and ranges for *J. frontalis* which are the black dots off the west coast of South America, and *J. caveorum* which is the black dot beneath the black arrow to the west of *J. frontalis*. Maps B and E show the habitat suitability results and range for *J. edwardsii* around both southern Australia and New Zealand. Maps C and F show the habitat suitability results and ranges for *J. lalandii* around South Africa, *J. paulensis* which are the black dots to the southeast of *J. lalandii*, and *J. tristani* which are the black dots to the southwest of *J. lalandii*.

## 3.3 Habitat modelling for climate scenario RCP60

The modelling results for climate scenario RCP60 were very similar to the RCP45 scenario. The percent of highly suitable locations were nearly identical to the RCP45 scenario (Table 4) with the same most important environmental variables (Table 3). The most notable change was for the *J. caveorum* habitat range which increased to 3% in highly suitable locations (Table 4). Also, there were very few changes to how the areas around *Jasus* habitat ranges appear under this scenario compared to RCP45 (Figures 5). Similar highly suitable locations were found along the west coast of South Africa, off the east coast of South America, and off both the west and east coasts of New Zealand for 2040-2050 and 2090-2100. Similar to the RCP45 scenario, simulations for RCP60 show large regions and small areas scattered outside of the current *Jasus* habitat range that contain slightly suitable conditions (coloured yellow or green) (Figures 5).

**Figure 5.**
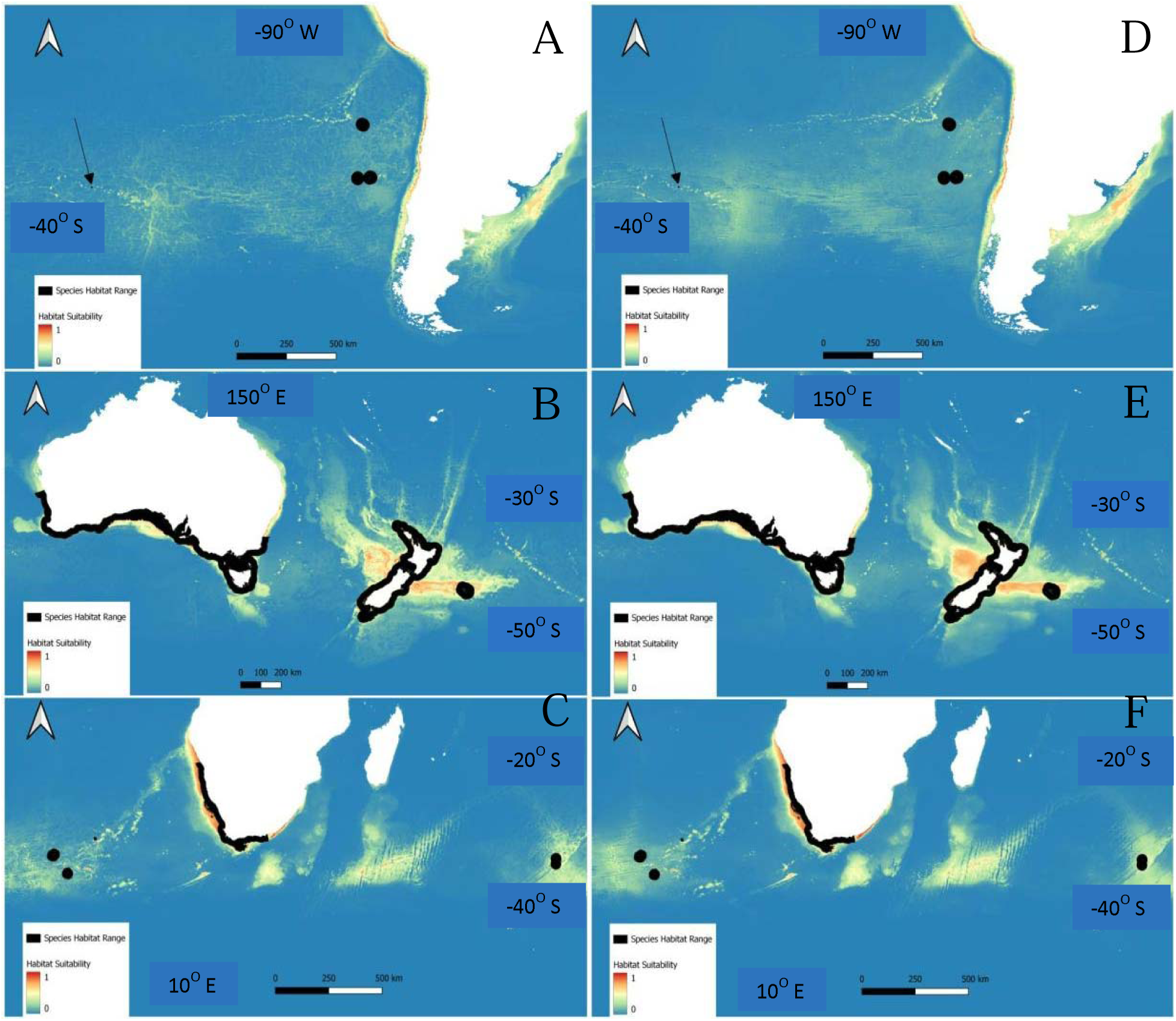
*Jasus* lobster MaxEnt habitat suitability maps for climate scenario RCP60 2040-2050 (A-C) and 2090-2100 (D-F) future periods. Maps A and D show the habitat suitability results and ranges for *J. frontalis* which are the black dots off the west coast of South America, and *J. caveorum* which is the black dot beneath the black arrow to the west of *J. frontalis*. Maps B and E show the habitat suitability results and range for *J. edwardsii* around both southern Australia and New Zealand. Maps C and F show the habitat suitability results and ranges for *J. lalandii* around South Africa, *J. paulensis* which are the black dots to the southeast of *J. lalandii*, and *J. tristani* which are the black dots to the southwest of *J. lalandii*.

## 3.4 Habitat modelling for climate scenario RCP85

Under climate scenario RCP85, all *Jasus* species will be subjected to the most dramatic environmental changes. While the most important variables did not change (Table 3), there was an overall decrease in highly suitable locations within species ranges compared to the previous two scenarios in the 2040-2050 time period (Table 4). Only *J. lalandii* showed an addition in the percent of highly suitable habitat in the RCP85 2040-2050 compared to the two previous climate scenarios increasing from 75% under RCP45 and RCP60 to 80% in RCP85. *J. paulensis* had the largest reduction in the percent of highly suitable locations decreasing from 71% under RCP45 and RCP60 to 27%. Under the RCP85 scenario, all species, except *J. caveorum,* had an increase in their percent of highly suitable habitat when comparing the 2040-2050 period to the 2090- 2100 period. The largest increase in the percent of highly suitable habitat was for *J. paulensis* going from 27% up to 71% (Table 4).

Modelling results for RCP85 showed similar trends compared to RCP45 and RCP60, but with a more southern shift in highly suitable habitat locations (Figures 4, 5). There was a small highly suitable area located farther south off the east coast of South America in 2040-2050 compared to previous results, and this area was predicted to become larger in 2090-2100 (Figure 6A, 6D). This was most evident off the west coast of New Zealand, off the east coast of southern South America, and just beyond the range of *J. lalandii* around South Africa (Figure 6).

**Figure 6.**
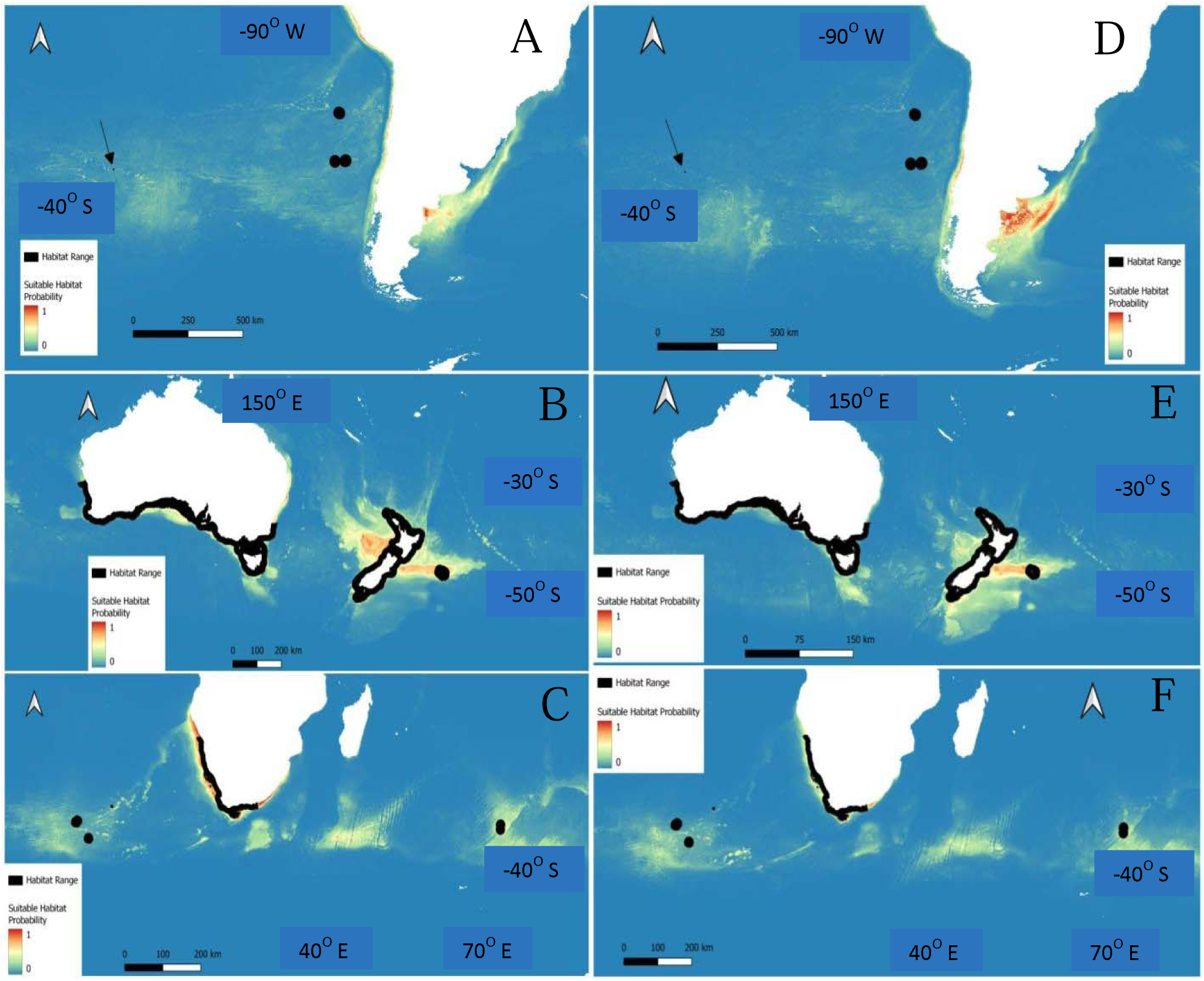
*Jasus* lobster MaxEnt habitat suitability maps for climate scenario RCP85 2040-2050 (A-C) and 2090-2100 (D-F) future periods. Maps A and D show the habitat suitability results and ranges for *J. frontalis* which are the black dots off the west coast of South America, and *J. caveorum* which is the black dot beneath the black arrow to the west of *J. frontalis*. Maps B and E show the habitat suitability results and range for *J. edwardsii* around both southern Australia and New Zealand. Maps C and F show the habitat suitability results and ranges for *J. lalandii* around South Africa, *J. paulensis* which are the black dots to the southeast of *J. lalandii*, and *J. tristani* which are the black dots to the southwest of *J. lalandii*.

### 4.0. Discussion

Benthic temperature was the most important environmental variable in determining suitable habitat locations for *Jasus* lobsters. For individual species, benthic temperature was one of the most important variables for *J. edwardsii* and *J. lalandii*, while the most important variables for *J. paulensis*, *J. frontalis*, and *J. tristani* were benthic nutrients (phosphate, silicate, and nitrate). For *J. caveorum,* there were several most important variables at both the surface and benthic layers. However, the low number of unique occurrence points and the small distribution range for *J. caveorum* created inaccurate modelling results. MaxEnt requires more than two unique occurrence points to train and test the model. Modelling results also found that the present habitat range for each lobster species differed in the percent of highly suitable locations. For the present period, those percentages ranged from the high 80s down to zero. For most species, MaxEnt results for the climate scenarios RCP45 and RCP60 predicted an increase in highly suitable locations compared to the present. Under scenario RCP85, species saw a reduction in percentages of those locations in 2040-2050, but saw an increase in 2090-2100.

Also, the limited number of environmental layers available for future projections could affect these estimates and results need to be interpreted with caution. Regardless, the overall southern movement of suitable habitat locations is highly likely as climate change continues to alter the marine environment.

## 4.1. Benthic temperature limiting lobster spatial distributions

This study provides evidence that the current habitat range of *Jasus* lobsters, specifically *J. edwardsii* and *J. lalandii*, are dependent on specific benthic temperatures. Water temperature limits the range of the different habitat types that marine lobsters occupy. *Jasus* lobsters preferentially inhabit kelp forest habitats which provide them with food, shelter, and are their preferred recruitment sites (Hinojosa et al. 2015). Kelp forests flourish in temperate regions, but increases in water temperature provide better conditions for algal turf (Provost et al. 2017). With changes to ocean temperatures, kelp forests are being dramatically reduced resulting in fewer preferred habitat locations for these lobsters (Hinojosa et al. 2015; Hesse et al. 2016). Therefore, lobster presence in new southern areas is dependent on kelp settlement and the establishment of healthy kelp forests within those areas. This limits the locations where lobsters can inhabit, and could reduce the number of regions where lobster fisheries can harvest from.

Different lobster life stages vary in their optimal thermal windows. For adult *J. lalandii*, the temperature range of 10-13°C enables normal activity levels, while any temperature increase or decrease by more than 5°C outside of that range caused elevated stress levels and decreased lobster activity (Zoutendyk 1989). For adult *J. edwardsii,* translocation of lobsters from deep to shallow waters resulted in faster growth rates compared to individuals inhabiting shallow and deep waters (Chandrapavan et al. 2010). Additionally, *J. edwardsii* juveniles thrive within a warmer thermal window of 19 to 21°C where they have optimal growth, survival, and feeding rates (Thomas et al. 2000). *Jasus* lobsters appear to grow faster in warmer water, and it is possible that rising ocean temperatures will initially accelerate their growth rates.

Comparatively, for the Eastern Rock lobster (*Sagmariasus verreauxi*) larvae, water temperature affects their development and survival with higher temperatures (ranging from 23- 25°C) favouring survival (Cetina-Heredia et al. 2015). Adult *S. verreauxi* lobsters are better suited to cooler temperatures of approximately 20°C (Cetina-Heredia et al. 2015) which is higher than adult *Jasus* lobsters. The California spiny lobster (*Panulirus interruptus*) larvae and adults have their highest densities in cold Arctic waters (Funes-Rodriguez et al. 2015). Their larvae and early life stages occur in higher abundances during warm months (summer-autumn), while the later life stages reach their highest abundances in colder months (winter-spring) (Funes-Rodriguez et al. 2015). However, larvae are most often found at the surface, while adults occur at deeper depths where temperatures are colder. Slight changes in water temperatures can create conditions that favour different life stages or species.

## 4.2. Other environmental variables influencing lobsters

While benthic temperature is one of the most important environmental variables for species located on a continental shelf, the most importance variables for species located around islands and on seamounts are benthic nutrients (nitrate, silicate, and phosphate) which are influenced by ocean pH. Altering ocean pH outside of lobsters’ optimal conditions will require them to allocate more energy toward survival instead of growth or reproduction (Knapp et al. 2016). Also, low pH levels decrease the availability of exoskeleton formation nutrients (such as calcium and carbon) which lobsters need to form and repair their shells (Hinojosa et al. 2017, 2018). This reduces the survival ability of lobsters as the exoskeleton provides protection, and low levels of readily available nutrients in the environment results in a thinner exoskeleton.

Ocean pH also limits the extents of the current habitat ranges for *J. edwardsii* and *J. lalandii*. The northern extent of the current range for *J. lalandii* is limited by an upwelling zone located just beyond the northern tip of its range (Jeffs et al. 2013), and for *J. edwardsii*, there is an upwelling area along the southern central coast of Australia (Linnane et al. 2010). Those upwelling regions alter the surrounding pH which limits the number of lobsters that are able to settle and survive there (Linnane et al. 2010; Jeffs et al. 2013). This suggests that pH is an important environmental variable for *Jasus* lobsters, especially the continental shelf inhabiting species that occupy areas influenced by a combination of factors (climate change, human activities, and upwelling zones).

Although ocean currents were not available and not modelled in this study, they play an important role in lobster distribution. As the larval phase lasts for up to two years in some species, lobsters have the potential to widely disperse and reach distant locations that have favourable habitat conditions (Bradford et al. 2015; Hinojosa et al. 2016; Chiswell & Booth 2017). Oceanic processes (such as fronts and currents) have the capacity to transport larvae over long distances and potentially between oceans. For example, the Subtropical Front, which connects the Atlantic to the Indian Ocean, is a pathway for larval dispersal between *J. tristani* and *J. paulensis* (Silva et al. 2021b). There is also evidence of gene flow in the opposite direction (i.e. from the Indian to the Atlantic Ocean) via the Agulhas leakage (Silva et al. 2021b). However, this east to west connectivity is expected to become weaker and result in a shift in the source-sink dynamics of these species as they move southwards tracking their optimal thermal conditions (Silva et al. 2021b).

As *Jasus* lobster habitat ranges are influenced by multiple environmental variables, there is a higher risk of profound impacts in lobster populations due to climate change. Changes to multiple environmental variables that were simultaneously affecting the habitats of the Indian spiny lobster (*Panulirus homarus*) have been previously shown to negatively influence their immune systems (Verghese et al. 2007). When there were more frequent environmental fluctuations, the lobsters became more susceptible to pathogens (Verghese et al. 2007). Having several simultaneously changing environmental variables might negatively impact *Jasus* lobsters through increasing their susceptibility to health changes. This likely results in higher lobster mortality, and will require a larger allocation of energy toward boosting their immune systems.

## 4.3. Implications of climate change for the lobster fishery

Commercial lobster fisheries will likely be impacted from changes in lobster settlement rates caused by climate change. For *J. edwardsii*, there were decreased juvenile lobster settlement rates around Tasmania, while those rates were at their highest for the states of South Australia and Victoria in 2005 and 2006 (Linnane et al. 2010). Those changes in juvenile settlement rates were attributed to large scale environmental factors and influenced the catches by local lobster fisheries over those years (Linnane et al. 2010). As a comparison, the tropical *Panulirus* lobster fishery in Indonesia was found to have decreased catches of *P. homarus* and *P. penicillatus* due to extended monsoon periods, but recovered when conditions returned to normal (Milton et al. 2014). These environmental changes created conditions that resulted in periods of altered catches for the fisheries and affected settlement rates. The combined effects of species southern range shifts and increased effects from climate change could result in dramatic changes to yearly catches for the lobster fisheries.

Regardless of whether the *Jasus* lobsters will be forced out of their current ranges or to more southern locations within them, these lobster fisheries already face difficulties as some are already considered to be fully exploited (Jeffs et al. 2013). The *J. edwardsii* fishery in the state of South Australia has decreased from 185 tonnes to 60 tonnes of lobsters caught per year from 1993 to 2005 (Linnane & Crosthwaite 2009). It is interesting to note that they had two to three times higher catch rates in offshore locations despite 80% of their catches coming from inshore areas (Linnane & Crosthwaite 2009). While some lobster fisheries are already facing difficulties, they might be able to counter this by moving with the lobsters to their new habitat locations.

The southern shift to new habitat locations for the *Jasus* lobsters could result in conflicts arising between neighbouring nation states or between countries. For *J. edwardsii,* all of the southern Australian states have commercial fisheries for this species (Linnane et al. 2010). The predicted southern shift in this study suggests that there will be a decrease in suitable habitat locations within the jurisdiction of the state of South Australia. This would cause reductions in catches there and possibly lead to a conflict arising with its neighbouring states of Victoria and Tasmania, as they are predicted to contain high concentrations of suitable habitat locations.

Issues could also arise for the countries currently harvesting *J. paulensis, J. frontalis,* and *J. tristani*. If these species move outside of each country’s exclusive economic zone, then conflicts could arise with other countries or companies that start harvesting lobsters in those new locations. These redistributions of species ranges need to be handled with an increased level of cooperation and governance between local, national, and international stakeholders.

### 4.4. Conclusion

In this study, benthic temperature was the most important environmental variable for the current spatial distribution of *Jasus* lobsters. Species located on a continental shelf were influenced the most by benthic temperature, while island and seamount dwelling species were more influenced by benthic nutrients. Modelling showed that the present habitat ranges of *Jasus* lobsters contained very different percentages of highly suitable locations. MaxEnt modelling identified other areas outside of their current habitat ranges that could be suitable. However, the model mainly predicted those locations to contain slightly suitable areas with some highly suitable locations scattered around. This suggests that the best suited habitat locations were located within their current habitat range. This study predicted that suitable habitat locations will move south as climate change alters ocean conditions. This movement will concentrate suitable habitats to the southern extents of their current range.

## Acknowledgements

Funding for this research was provided by an Australian Research Council Discovery Project grant Project No. DP150101491.

## Notes

### Competing Interest Statement

The authors have declared no competing interest.

### Summary of Updates

Supplemental files updated.

