## Supplemental Figures for "Environmental determinants of suitable habitat and the prediction of a southern shift in the future distribution of spiny lobsters, genus *Jasus*"

**Supplementary Material**

J. caveorum

Percentage

| Climate Scenario | Not Suitable | Slightly | Moderately | Highly |
| --- | --- | --- | --- | --- |
| Present | 1 | 0 | 0 | 0 |
| RCP45 2040-2050 | 1 | 0 | 0 | 0 |
| RCP45 2090-2100 | 0.967742 | 0.0322581 | 0 | 0 |
| RCP60 2040-2050 | 0.967742 | 0.0322581 | 0 | 0 |
| RCP60 2090-2100 | 0.942857 | 0.0285714 | 0 | 0.0285714 |
| RCP85 2040-2050 | 1 | 0 | 0 | 0 |
| RCP85 2090-2100 | 1 | 0 | 0 | 0 |

J. edwardsii

Percentage

| Climate Scenario | Not Suitable | Slightly | Moderately | Highly |
| --- | --- | --- | --- | --- |
| Present | 0.000946882 | 0.00466479 | 0.122255 | 0.872134 |
| RCP45 2040-2050 | 0.00329088 | 0.0128757 | 0.0629862 | 0.920847 |
| RCP45 2090-2100 | 0.00175916 | 0.00468799 | 0.0534617 | 0.940091 |
| RCP60 2040-2050 | 0.00315627 | 0.0116875 | 0.0535174 | 0.931639 |
| RCP60 2090-2100 | 0.00222332 | 0.0042424 | 0.0578062 | 0.935728 |
| RCP85 2040-2050 | 0.00150387 | 0.00502683 | 0.21659 | 0.776879 |
| RCP85 2090-2100 | 0.00201909 | 0.00891183 | 0.155934 | 0.833135 |

J. frontalis

Percentage

| Climate Scenario | Not Suitable | Slightly | Moderately | Highly |
| --- | --- | --- | --- | --- |
| Present | 0.230769 | 0 | 0.230769 | 0.538462 |
| RCP45 2040-2050 | 0 | 0 | 0.769231 | 0.230769 |
| RCP45 2090-2100 | 0 | 0 | 0.769231 | 0.230769 |
| RCP60 2040-2050 | 0 | 0 | 0.769231 | 0.230769 |
| RCP60 2090-2100 | 0 | 0 | 0.769231 | 0.230769 |
| RCP85 2040-2050 | 0 | 0 | 0.846154 | 0.153846 |
| RCP85 2090-2100 | 0 | 0 | 0.692308 | 0.307692 |

J. lalandii

Percentage

| Climate Scenario | Not Suitable | Slightly | Moderately | Highly |
| --- | --- | --- | --- | --- |
| Present | 0.0203046 | 0.0126904 | 0.312183 | 0.654822 |
| RCP45 2040-2050 | 0.0329949 | 0.0558376 | 0.167513 | 0.743655 |
| RCP45 2090-2100 | 0.0329949 | 0.035533 | 0.137056 | 0.794416 |
| RCP60 2040-2050 | 0.0329949 | 0.0609137 | 0.164975 | 0.741117 |
| RCP60 2090-2100 | 0.0304569 | 0.0380711 | 0.109137 | 0.822335 |
| RCP85 2040-2050 | 0.0126904 | 0.0253807 | 0.119289 | 0.84264 |
| RCP85 2090-2100 | 0.0101523 | 0.0279188 | 0.0609137 | 0.901015 |

J. paulensis

Percentage

| Climate Scenario | Not Suitable | Slightly | Moderately | Highly |
| --- | --- | --- | --- | --- |
| Present | 0.272727 | 0 | 0.272727 | 0.454545 |
| RCP45 2040-2050 | 0.272727 | 0 | 0 | 0.727273 |
| RCP45 2090-2100 | 0.272727 | 0 | 0.0909091 | 0.636364 |
| RCP60 2040-2050 | 0.272727 | 0 | 0 | 0.727273 |
| RCP60 2090-2100 | 0.272727 | 0 | 0.0909091 | 0.636364 |
| RCP85 2040-2050 | 0.272727 | 0 | 0.454545 | 0.272727 |
| RCP85 2090-2100 | 0.272727 | 0 | 0 | 0.727273 |

J. tristani

Percentage

| Climate Scenario | Not Suitable | Slightly | Moderately | Highly |
| --- | --- | --- | --- | --- |
| Present | 0.444444 | 0.0555556 | 0.5 | 0 |
| RCP45 2040-2050 | 0.222222 | 0.0555556 | 0.222222 | 0.5 |
| RCP45 2090-2100 | 0.222222 | 0 | 0.277778 | 0.5 |
| RCP60 2040-2050 | 0.222222 | 0 | 0.277778 | 0.5 |
| RCP60 2090-2100 | 0.222222 | 0 | 0.166667 | 0.611111 |
| RCP85 2040-2050 | 0.222222 | 0.166667 | 0.277778 | 0.333333 |
| RCP85 2090-2100 | 0.222222 | 0.166667 | 0.222222 | 0.388889 |

Jasus

Percentage

| Climate Scenario | Not Suitable | Slightly | Moderately | Highly |
| --- | --- | --- | --- | --- |
| Present | 0.00119031 | 0.00468248 | 0.122629 | 0.871498 |
| RCP45 2040-2050 | 0.00351997 | 0.0129544 | 0.0632205 | 0.920305 |
| RCP45 2090-2100 | 0.00198693 | 0.00474733 | 0.0536703 | 0.939595 |
| RCP60 2040-2050 | 0.00338102 | 0.011778 | 0.0537722 | 0.931069 |
| RCP60 2090-2100 | 0.00244545 | 0.00430733 | 0.0579452 | 0.935302 |
| RCP85 2040-2050 | 0.00169977 | 0.00507617 | 0.216436 | 0.776788 |
| RCP85 2090-2100 | 0.00220924 | 0.00895739 | 0.155768 | 0.833065 |

Figure 1 Percentage of suitable habitat locations within the current habitat range of A) *J. caveorum,* B) *J. edwardsii*, C) *J. frontalis*, D) *J. lalandii*, E) *J. paulensis*, F) *J. tristani*, and G) genus *Jasus* lobsters from the present period (2000-2014) and two future periods (2040-2050 and 2090-2100) for climate scenarios RCP45, RCP60, and RCP85.

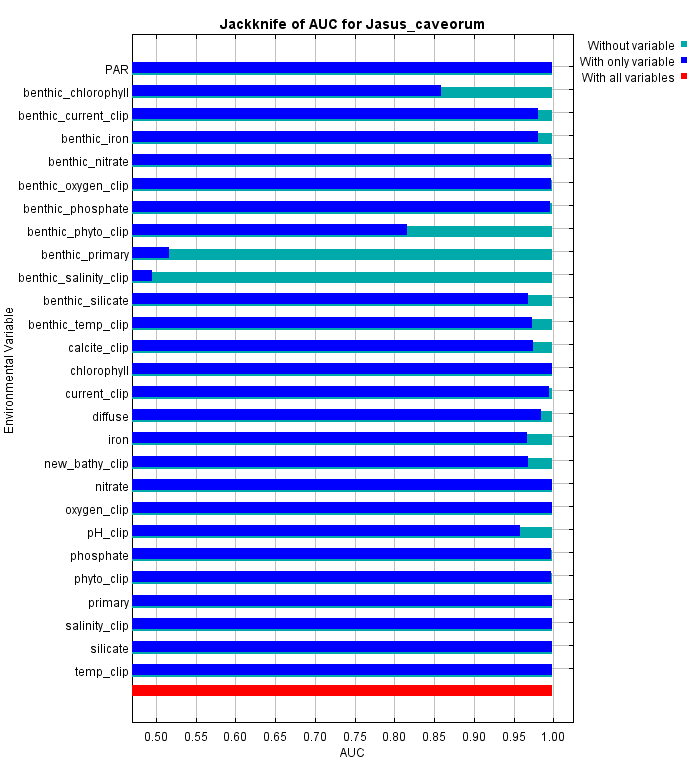

A

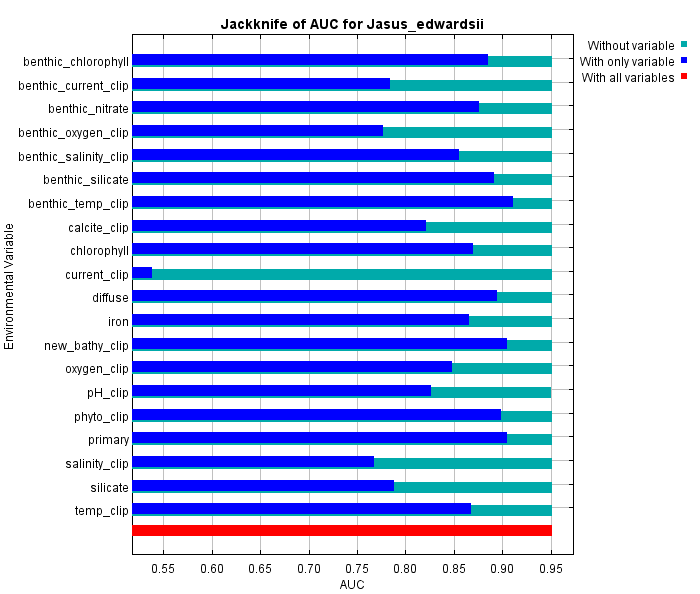

B

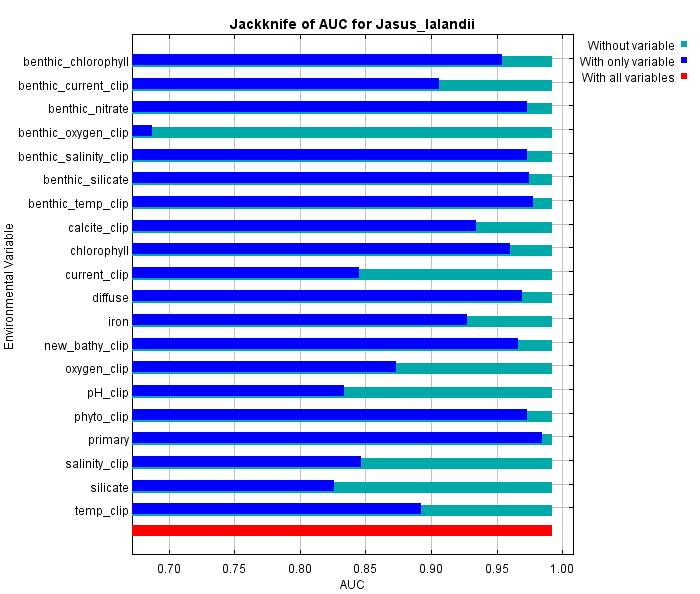

C

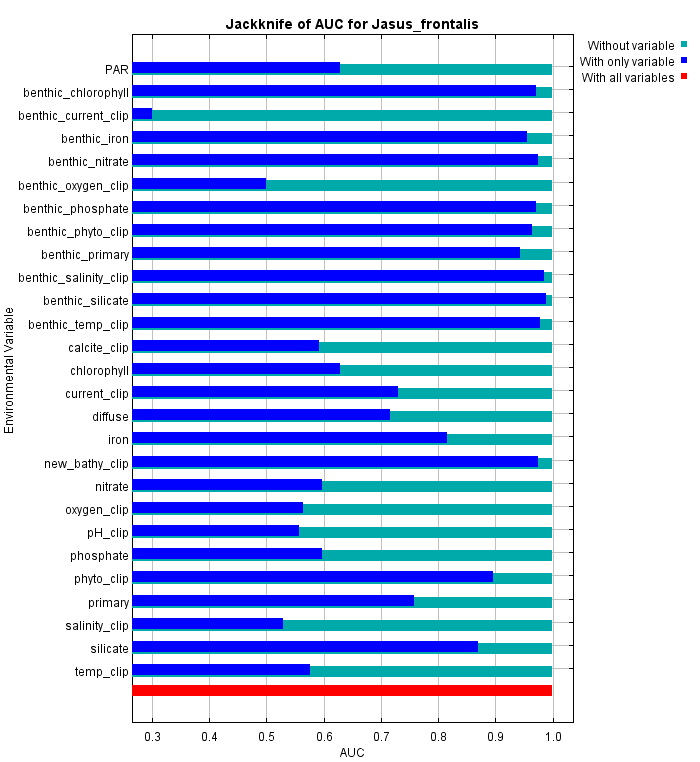

D

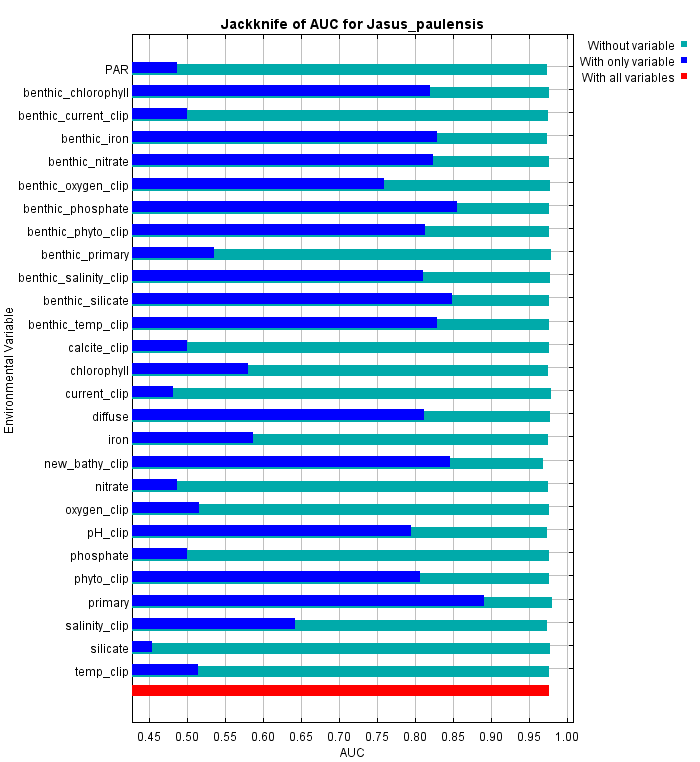

E

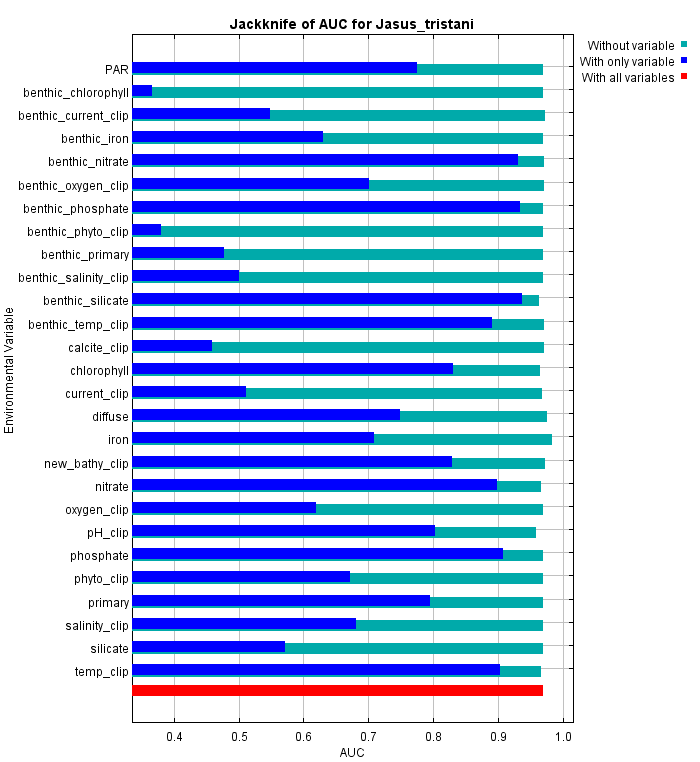

F

Figure 2 Area under the curve (AUC) jackknife results from MaxEnt simulations for A) *J. caveorum*, B) *J. edwardsii*, C) *J. lalandii*, D) *J. frontalis*, E) *J. paulensis*, and F) *J. tristani* for the present (2000-2014) indicating the importance of each environmental variable to the model when using only that variable (dark blue) or when excluding that variable (light blue).

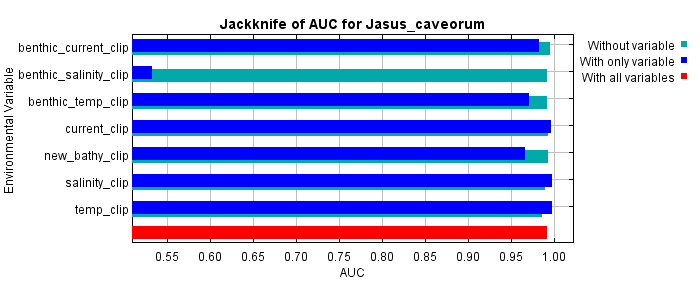

A

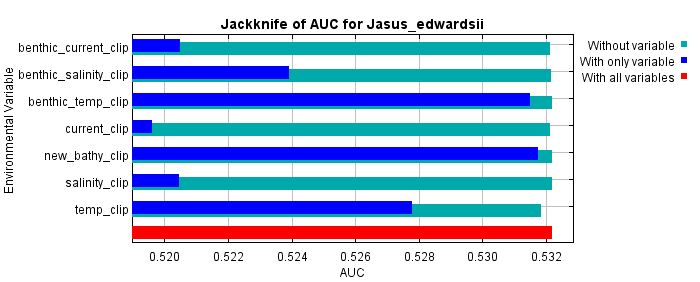

B

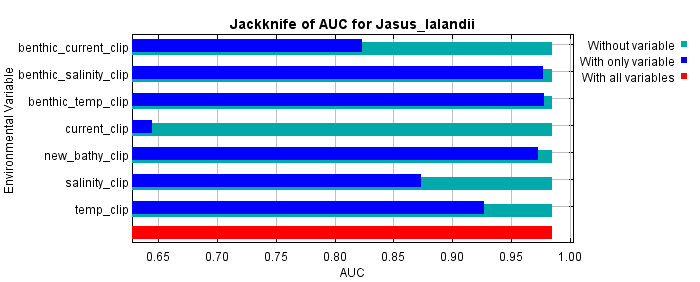

C

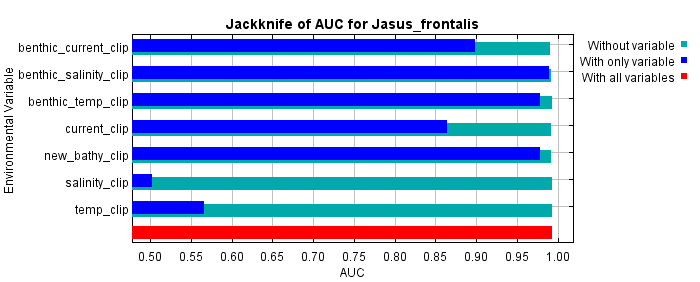

D

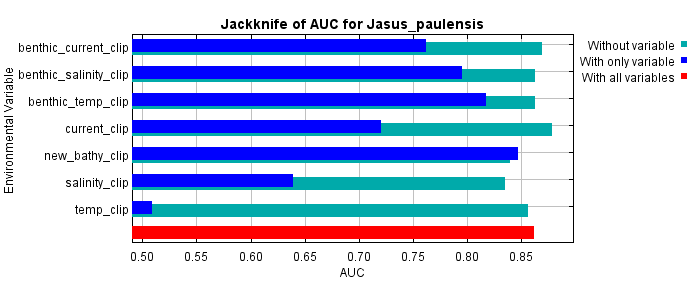

E

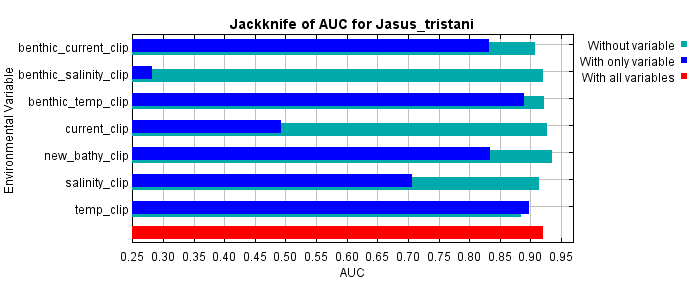

F

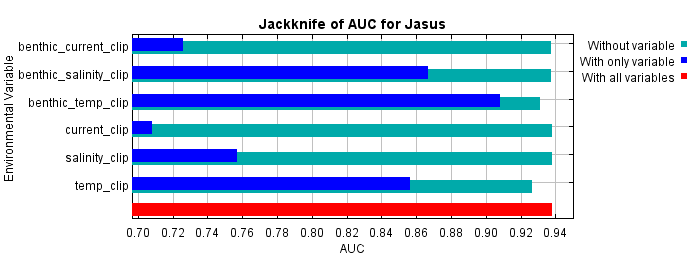

G

Figure 3 Area under the curve (AUC) jackknife results from MaxEnt simulations for A) *J. caveorum*, B) *J. edwardsii*, C) *J. lalandii*, D) *J. frontalis*, E) *J. paulensis*, F) *J. tristani*, and G) genus *Jasus* for the RCP45 scenario in 2040-2050 indicating the importance of each environmental variable to the model when using only that variable (dark blue) or when excluding that variable (light blue).

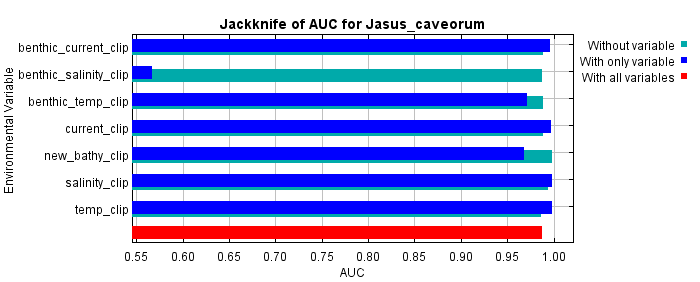

A

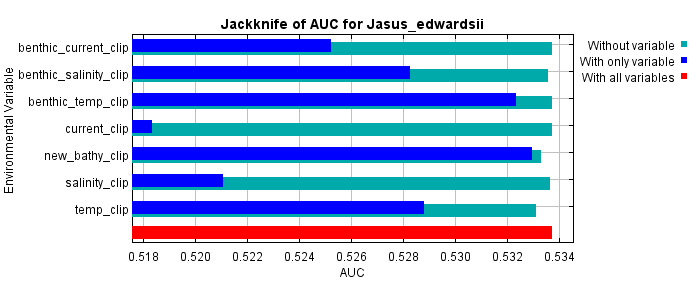

B

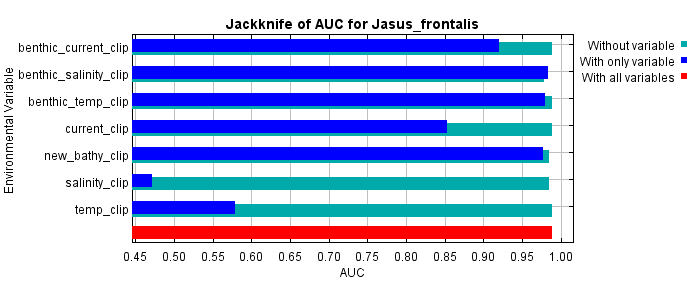

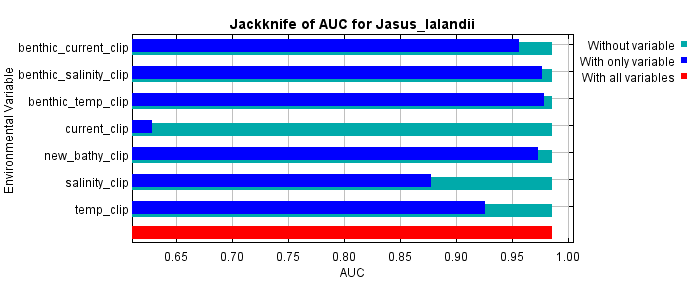

D

C

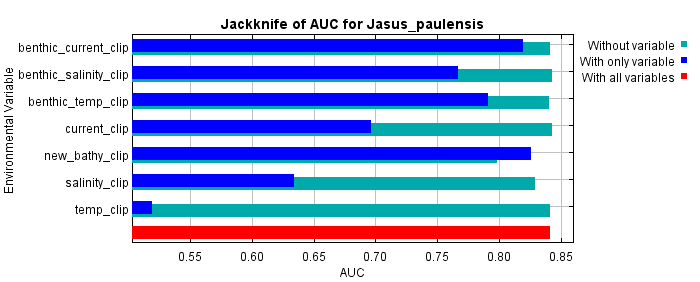

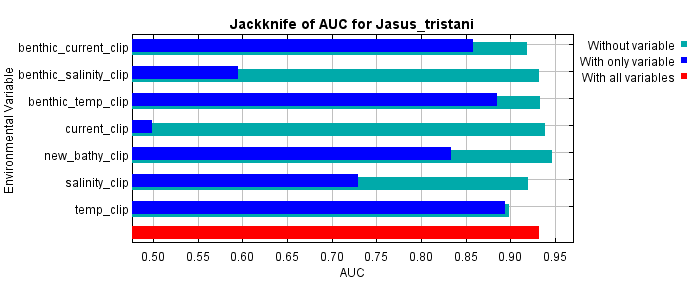

F

E

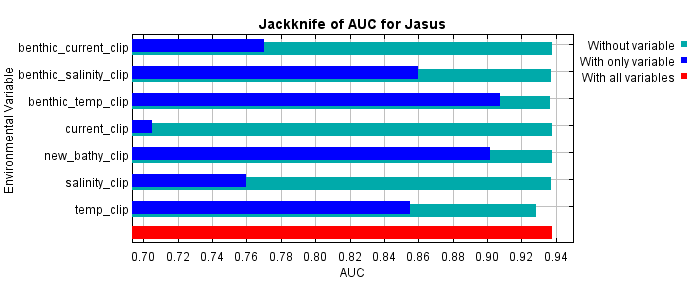

G

Figure 4 Area under the curve (AUC) jackknife results from MaxEnt simulations for A) *J. caveorum*, B) *J. edwardsii*, C) *J. lalandii*, D) *J. frontalis*, E) *J. paulensis*, F) *J. tristani,* and G) genus *Jasus* for the RCP45 scenario in 2090-2100 indicating the importance of each environmental variable to the model when using only that variable (dark blue) or when excluding that variable (light blue).

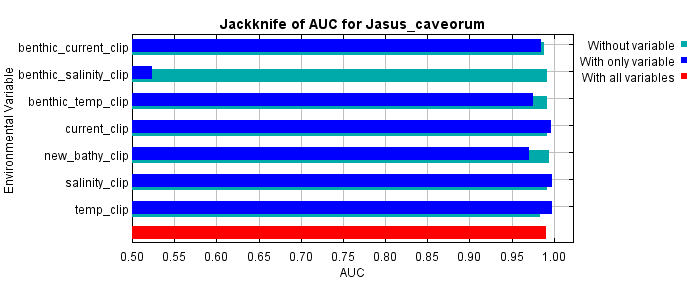

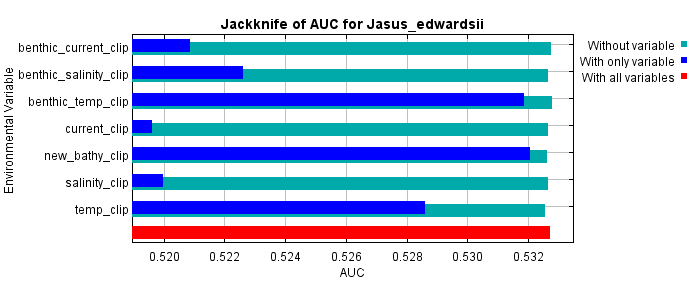

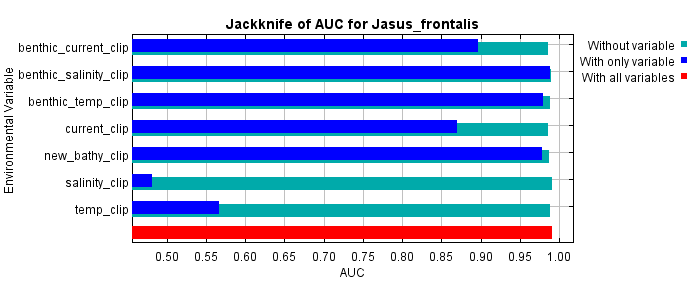

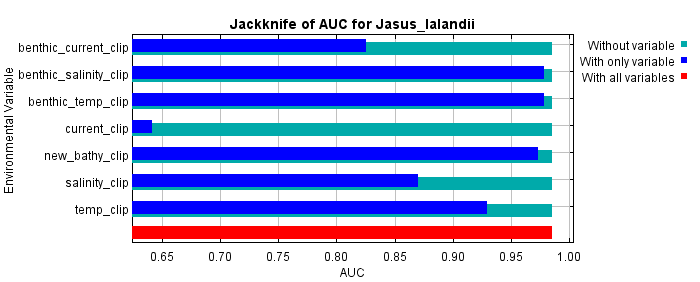

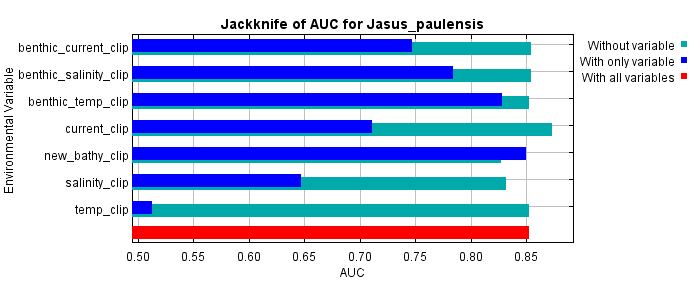

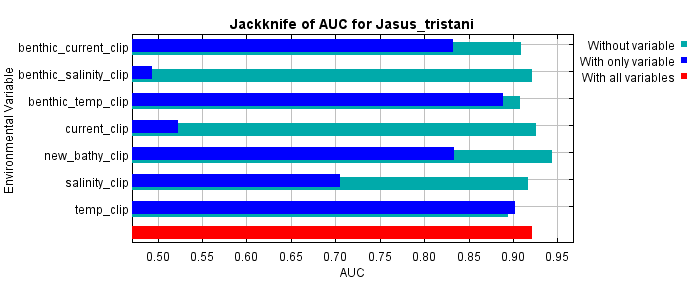

C

B

A

D

E

F

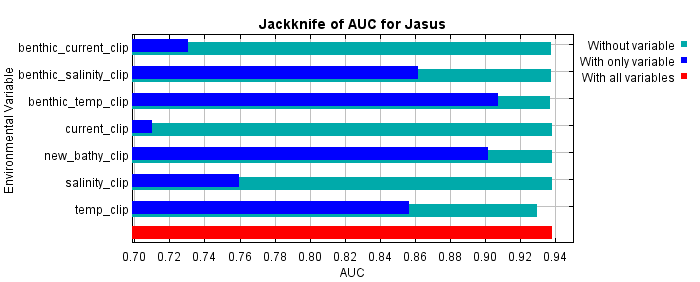

G

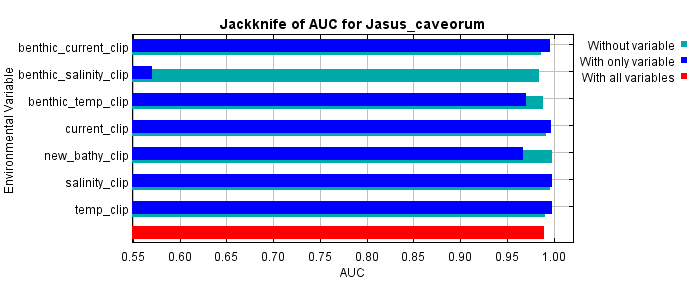
Figure 5 Area under the curve (AUC) jackknife results from MaxEnt simulations for A) *J. caveorum*, B) *J. edwardsii*, C) *J. frontalis*, D) *J. lalandii*, E) *J. paulensis*, F) *J. tristani,* and G) genus *Jasus* for the RCP60 scenario in 2040-2050 indicating the importance of each environmental variable to the model when using only that variable (dark blue) or when excluding that variable (light blue).

A

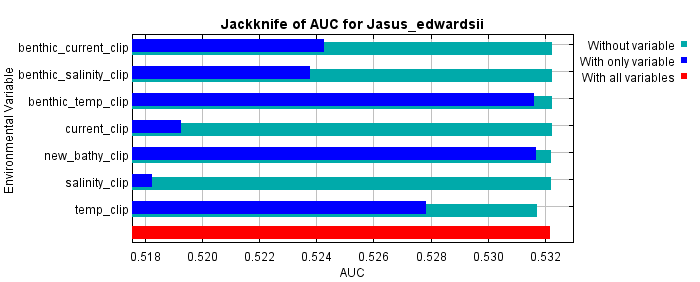

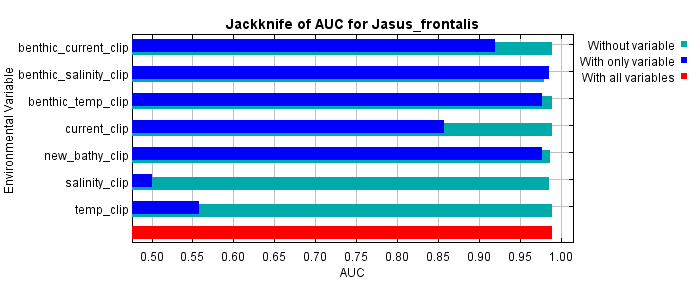

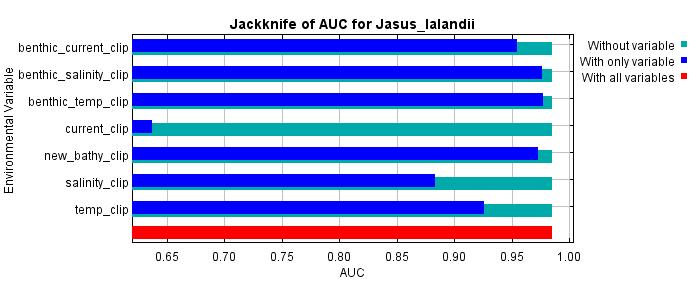

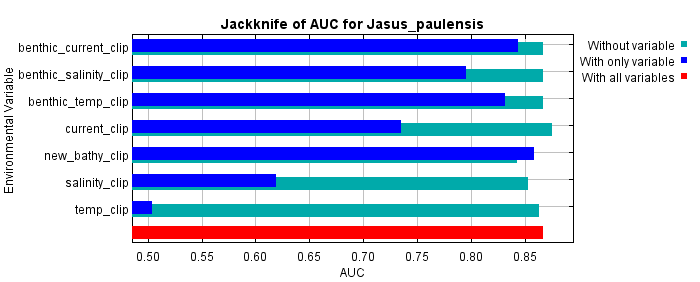

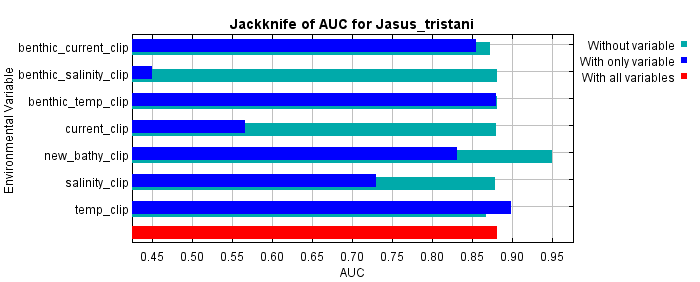

D

C

B

E

F

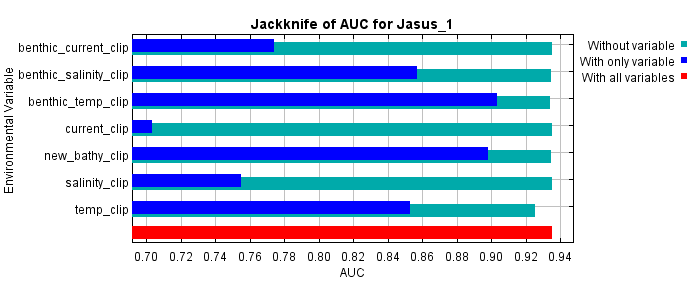

G

Figure 6 Area under the curve (AUC) jackknife results from MaxEnt simulations for A) *J. caveorum*, B) *J. edwardsii*, C) *J. frontalis*, D) *J. lalandii*, E) *J. paulensis*, F) *J. tristani,* and G) genus *Jasus* for the RCP60 scenario in 2090-2100 indicating the importance of each environmental variable to the model when using only that variable (dark blue) or when excluding that variable (light blue).

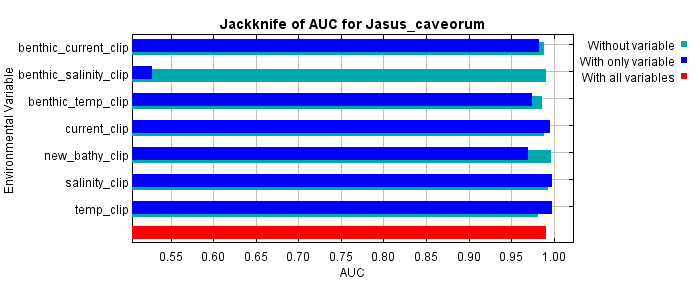

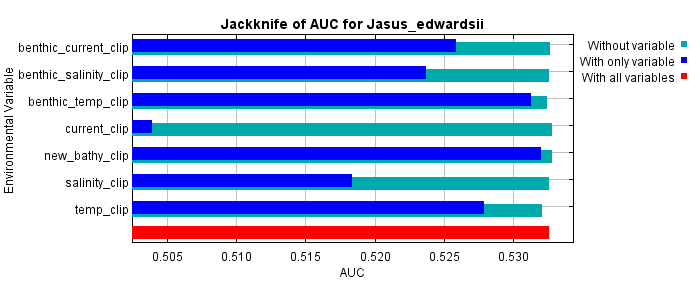

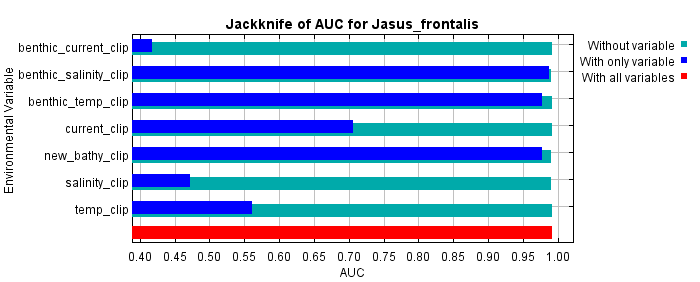

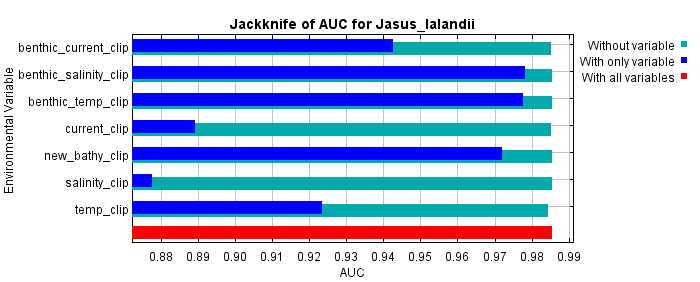

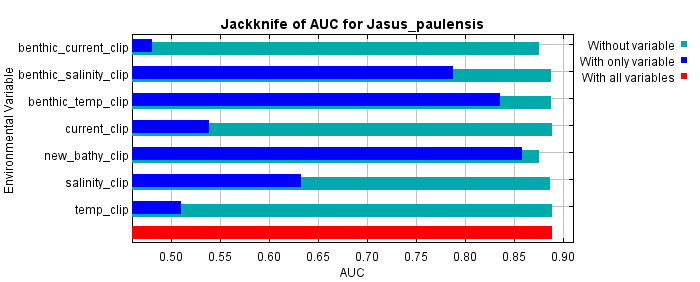

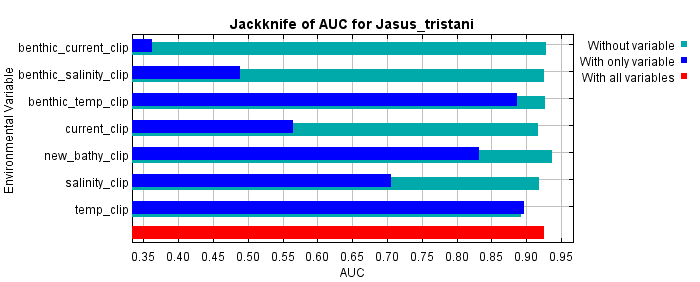

C

B

A

D

E

F

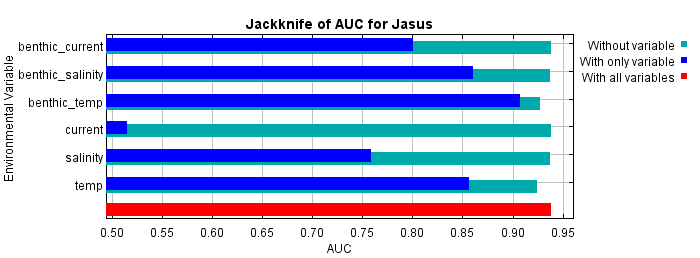

G

Figure 7 Area under the curve (AUC) jackknife results from MaxEnt simulations for A) *J. caveorum*, B) *J. edwardsii*, C) *J. frontalis*, D) *J. lalandii*, E) *J. paulensis*, F) *J. tristani*, and G) genus *Jasus* for the RCP85 scenario in 2040-2050 indicating the importance of each environmental variable to the model when using only that variable (dark blue) or when excluding that variable (light blue).

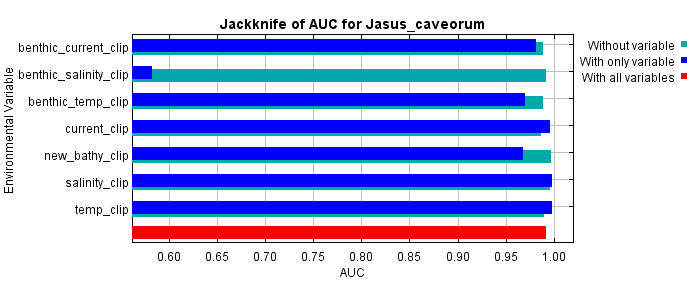

A

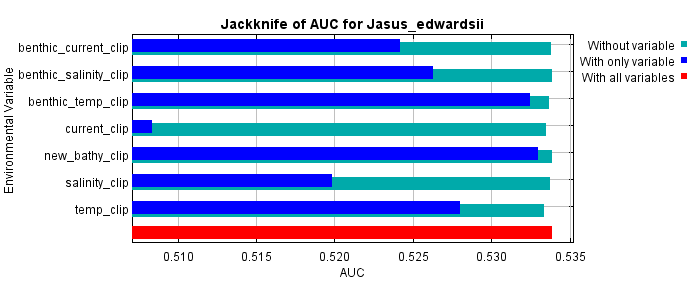

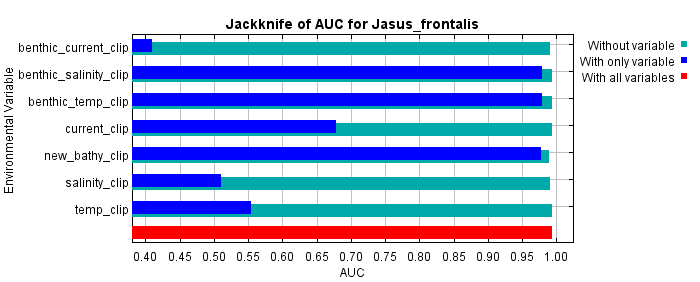

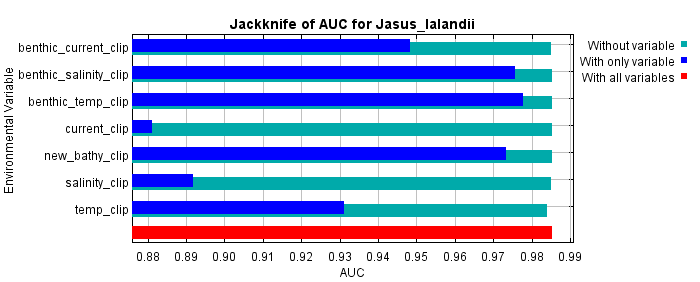

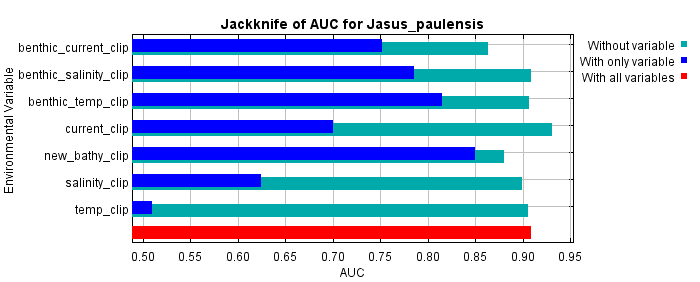

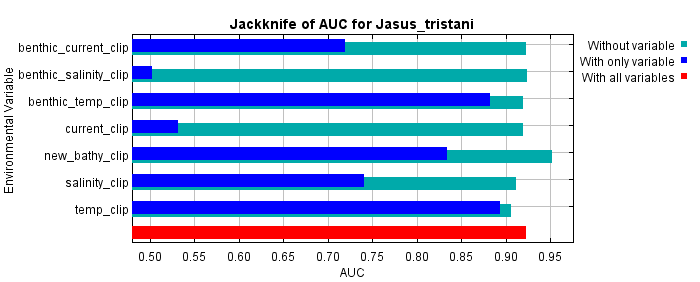

D

C

B

E

F

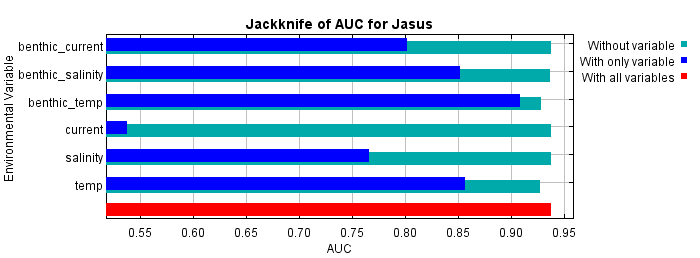

G

Figure 8 Area under the curve (AUC) jackknife results from MaxEnt simulations for A) *J. caveorum*, B) *J. edwardsii*, C) *J. frontalis*, D) *J. lalandii*, E) *J. paulensis*, F) *J. tristani*, and G) genus *Jasus* for the RCP85 scenario in 2090-2100 indicating the importance of each environmental variable to the model when using only that variable (dark blue) or when excluding that variable (light blue).
